# Nanoscale organization of the endogenous ASC speck

**DOI:** 10.1101/2021.09.17.460822

**Authors:** Ivo M. Glück, Grusha Primal Mathias, Sebastian Strauss, Thomas S. Ebert, Che Stafford, Ganesh Agam, Suliana Manley, Veit Hornung, Ralf Jungmann, Christian Sieben, Don C. Lamb

## Abstract

The NLRP3 inflammasome is a central component of the innate immune system. Its activation leads to the formation of a supramolecular assembly of the inflammasome adaptor protein ASC, commonly referred to as the “ASC speck”. Different models of the overall structure of the ASC speck, as well as the entire NLRP3 inflammasome, have been reported in the literature. While many experiments involve overexpression or *in vitro* reconstitution of recombinant ASC, the cytoplasmic endogenous ASC speck remains difficult to study due to its relatively small size and structural variability.

Here, we use a combination of fluorescence imaging techniques including dual-color 3D super-resolution imaging (dSTORM and DNA-PAINT) to visualize the endogenous ASC speck following NLRP3 inflammasome activation. We observe that the complex varies in diameter between ∼800 and 1000 nm and is composed of a dense core from which filaments extend into the periphery. We used a combination of anti-ASC antibodies as well as much smaller nanobodies for labeling, and find that the larger antibodies reliably label the lower-density periphery whereas the nanobody, which has a lower binding affinity, is more efficient in labeling the dense core. Imaging whole cells using dSTORM, furthermore, allowed us to sort the imaged structures into a pseudo-time sequence suggesting that the endogenous ASC speck becomes denser but not much larger during its formation. The reported results add an important piece of information towards a comprehensive understanding of the supramolecular structure of the endogenous inflammasome complex.

**Significance:** The assembly of ASC into a supramolecular complex referred to as the ASC speck is a critical step in the NLRP3 inflammasome response. Despite significance research, the organization and formation of the endogenous ASC speck remain elusive. We used a combination of differently sized ASC labels with super-resolution microscopy to address the organization of the ASC speck at the nanoscale. Our results allow inference of the supramolecular structure of the unperturbed, endogenous speck and characterize how it changes during ASC recruitment. We find that the complex varies in size and is composed of a dense core from which filaments extend into the periphery. Notably, our results further indicate that during inflammasome formation, ASC specks become denser but only slightly larger.

## Introduction

Inflammasomes are a class of large, multiprotein complexes that assemble upon activation of cellular pattern recognition receptors (PRRs) ^1^. Being part of the innate immune system, inflammasomes can sense the presence of non-self biomolecules or the perturbation of cellular homeostasis. The key components include a sensor protein, the adaptor protein Apoptosis-associated speck-like protein containing a Caspase activation and recruitment domain (ASC) ^2, 3^ and the inflammatory caspase 1 (CASP1) ^4–6^. The largest group of sensor proteins belongs to the family of nucleotide-binding oligomerization domain-like receptors (NLRs). Among these, NLRP3 (NOD-, LRR- and pyrin domain-containing protein 3) has been shown to play a critical role in many infectious, as well as sterile inflammatory conditions. Although the molecular mode of action remains to be elucidated, potassium efflux, as it occurs in the context of membrane damage, appears to be a key signal upstream of NLRP3 activation ^7^. NLRP3 activation triggers recruitment of ASC and CASP1 leading to a single micrometer-sized assembly. For the structure of ASC within the inflammasome, Masumoto *et al.* coined the term “ASC speck” ^3, 8^. ASC is composed of two interaction domains connected by a semi-flexible linker: a Pyrin domain (PYD) ^9^ and a Caspase activation and recruitment domain (CARD) ^10, 11^. The individual domains have a high tendency for homotypic interactions due to their complementarity in structure and charge, leading to the assembly of the large multiprotein inflammasome complex ^12–15^.

The details of how the ASC speck is organized is the subject of intensive research. ASC (22 kDa) is soluble at low pH and in a chaotropic solution, but at physiological pH, the protein can assemble into filaments as observed *in vitro* by solid-state NMR spectroscopy and electron microscopy (EM) ^16–21^. The ASC speck can also be purified from inflammasome-activated cells expressing ASC endogenously, where it was found to take on diverse morphologies, including a star-shaped assembly ^8^, isolated filaments, or amorphous aggregates potentially composed of intertwined filaments ^19, 22^. In cells, ASC specks were visualized by microscopy using immunofluorescence or expression of ASC tagged with a fluorescent protein. Upon overexpression, the resulting speck was typically much larger than 1 µm and occasionally showed filaments protruding from the edge of the structure ^20, 22–25^. In contrast, diffraction-limited immunofluorescence imaging of the endogenous ASC speck revealed a spot of about 1 µm in diameter. Due to the propensity of ASC to self-assemble, it is unclear whether the endogenous structure resembles the complexes observed *in vitro* or upon ASC overexpression. Interestingly, one approach to labeling ASC using a EGFP-labeled nanobody directed against the CARD domain revealed an intermediate, filamentous structure during speck formation, but no ASC specks were observed ^26^. Higher resolution *in situ* studies, either by EM ^27, 28^ or super-resolution fluorescence STED (stimulated emission depletion) microscopy ^29^ resolved the endogenous speck as an amorphous aggregate potentially made up of intertwined filaments. In contrast, other studies describe the ASC speck as a hollow, ring-shaped complex ^3, 6, 23, 30–38^, and based on this observation, different models for ASC speck and inflammasome formation have been proposed ^34, 35, 39–42^. Hence, despite its relevance for understanding inflammasome formation, the nanoscale organization of the endogenous ASC speck remains controversial.

Here, we performed a systematic study of fluorescence labelling strategies, and used quantitative widefield microscopy, confocal microscopy and single-molecule localization microscopy (i.e. 3D dual-color dSTORM and 3D DNA-PAINT) to investigate the organization of the endogenous ASC speck. Our data resolved filaments protruding from a dense core of the endogenous ASC speck, supporting the amorphous nature of the complex. By using two complementary labelling approaches comparing antibody-with nanobody-labelled ASC, we could probe the organization of and density differences within the ASC speck. We found that nanobodies labelled the center of the speck while the antibody was predominantly detected in the periphery of the complex, occasionally exhibiting a hollow center. Thus, our results reconcile the disparate structures reported in the literature. Furthermore, we analyzed the redistribution of ASC into the speck using single-cell analysis, allowing us to sort specks with respect to the degree of ASC recruitment. Our results indicate that endogenous specks mainly become denser but only slightly larger during inflammasome assembly.

## Results

### Endogenous specks vary strongly in ASC content

We investigated the organization of the endogenous ASC speck in THP-1 cells. The cells were primed using lipopolysaccharide (LPS) ^43^ followed by stimulation with the bacterial, potassium-efflux-inducing ionophore nigericin ^44^. After 90 minutes, the cells were fixed using paraformaldehyde and stained with a primary monoclonal antibody against ASC in combination with a secondary Alexa Fluor 647-tagged F(ab’)_2_ fragment. To gain an initial insight into the distribution of ASC in the cell, we imaged the cells at low magnification (10x) using confocal microscopy. In unstimulated cells, ASC was distributed throughout the cell, including the nucleus (**Figure 1A**, left panel, **Supplementary Figure S1A**). Upon NLRP3 inflammasome activation, ASC relocated into the characteristic single perinuclear speck in about 30% of the cells (**Figure 1A**, right panel) ^45^. We could also observe extracellular specks as previously reported ^22, 27^ (**Supplementary Figure S1B**). Next, we used high magnification (60x) widefield microscopy to image 59 individual ASC specks, which appear as spherical complexes ranging in size between approximately 0.5 - 1 µm diameter (**Figure 1B**). Analysis of the integrated intensity revealed that the amount of incorporated ASC as reflected by antibody binding can vary by almost one order of magnitude (**Figure 1C**). The widefield microscopy images also confirmed that ASC is distributed throughout unstimulated cells (**Figure 2A**, left panel), and that, in cells showing a speck, the cytoplasmic ASC was almost completely redistributed into a single, bright ASC spot (**Figure 2A**, right panel). These observations are consistent with previous reports of the ASC speck upon activation of the NLRP3 inflammasome ^46^.

**Figure 1.**
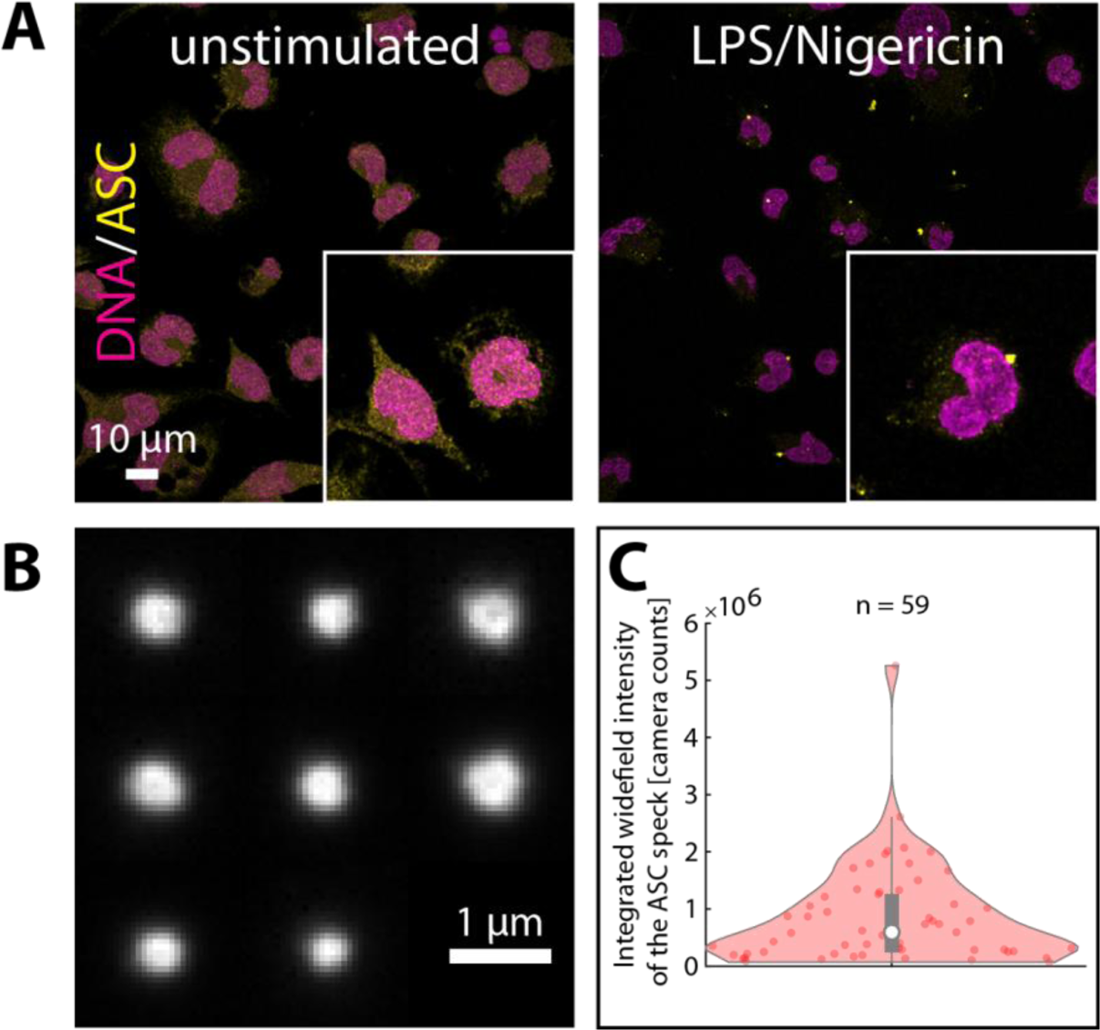
LPS + Nigericin stimulation of THP-1 cells induces ASC speck formation with variable size and ASC content. A) Confocal images (maximum projections) of fluorescently labeled ASC (shown in yellow) in unstimulated (left panel) and in LPS + Nigericin-stimulated THP-1 cells (right panel). The formation of ASC specks after stimulation is clearly visible. DNA was stained using DAPI (shown in magenta). B) A montage of ASC specks immunostained using a primary antibody and a secondary Alexa Fluor 647-conjugated F(ab’)_2_ fragment and imaged by diffraction-limited widefield illumination with 60x magnification. The speck size varies and the appearance of a darker center is observed for some of the specks. C) A violin plot of the integrated widefield intensity of ASC specks imaged as described for panel B. There is a large variation in the intensity, which serves as an indicator for the amount of protein incorporated into the speck. Pooled data from three independent cell preparations is shown.

**Figure 2.**
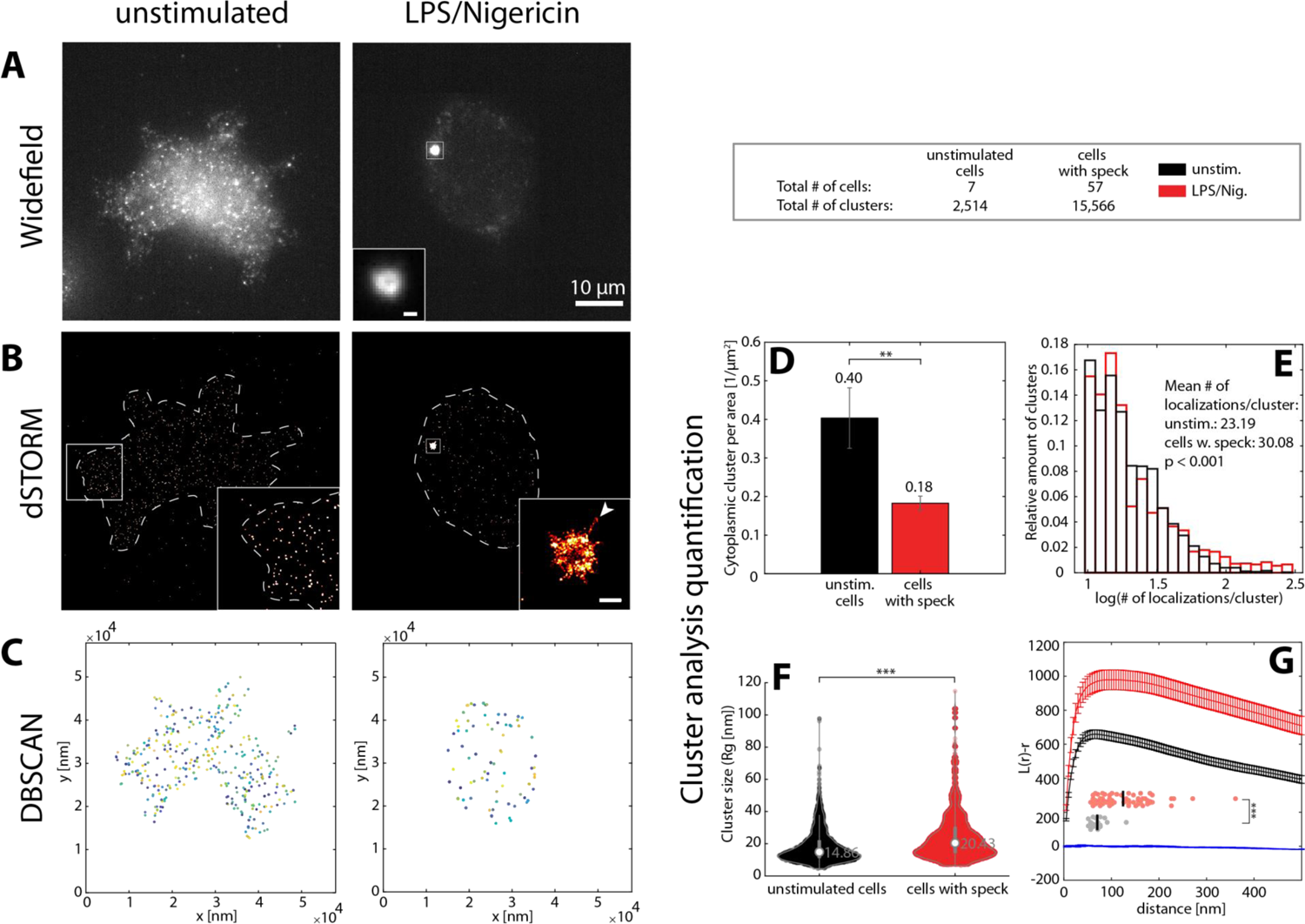
Whole-cell dSTORM super-resolution microscopy of endogenous ASC in THP-1 cells. A, B) The distribution of ASC in non-stimulated cells (left panels) and the redistribution of ASC into the speck after stimulation with LPS and Nigericin (right panels) observed using diffraction-limited widefield imaging (A) and dSTORM (B). dSTORM additionally resolves filaments protruding from the speck core (one example is highlighted by the arrow). Scale bars in insets correspond to 500 nm. C) DBSCAN clustering of (non-speck bound) cytosolic ASC localizations confirms depletion of the protein from the cytosol after speck formation. Color coding is used to distinguish individual clusters. D – G) The results of the DBSCAN analysis are shown. D) The non-speck incorporated ASC cluster density decreases in cells exhibiting a speck due to recruitment of the protein into the speck. The error bars indicate the standard error of the mean calculated from the analysis results of individual cells. E), F) The distribution of the number of localizations (E) and violin plots of the radius of gyration (*R_g_*) (F) for unstimulated cells and cells forming a speck. The distributions are similar with an additional population observable in speck-producing cells with more localizations and a larger size compared to non-stimulated cells. White circles in the violin plots indicate the median of the distribution. The statistical significance in D-F was assessed by a Two-Sample Kolmogorov-Smirnov Test (*) p<0.05, (**) p<0.01, (***) p<0.001. G) A Ripley’s K analysis of the ASC clusters. The Ripley’s K function confirms the decrease in cluster density after speck formation as well as the increase in cluster size. The curves show the mean ± SD of 100 ROI from 40 cells imaged from three independent experiments. For the estimation of the curve’s maximum (G, lower part), we measured from the individual ROIs). The same data was taken and the positions of the individual localizations were randomized at the same density and analyzed for comparison. The statistical significance was assessed by one-sided t-test (**) p<0.005, (***) p<0.001. Data on unstimulated cells was obtained on a single experiment. Data on cells with speck was obtained on three independent cell preparations.

### Non-speck incorporated ASC clusters maintain a similar size distribution during ASC speck formation

To study the distribution of ASC at the nanoscale, we used direct stochastic optical reconstruction microscopy (dSTORM) ^47^. Reconstructed dSTORM images further confirmed the cytoplasmic ASC distribution in unstimulated cells and its redistribution into a single speck after inflammasome activation. Interestingly, the increased detection sensitivity of dSTORM allowed us to visualize a previously undetected non-speck-bound ASC population in the cytoplasm of activated cells (**Figure 2B**). We performed a clustering analysis based on the local density of molecules, which distinguishes cytoplasmic ASC clusters from background localizations (**Supplementary Figure S2 and Supplementary Figure S3**). We used a Density-Based Spatial Clustering of Applications with Noise (DBSCAN) analysis ^48^ (**Figure 2C**), which revealed a decrease in the cluster density from an average of 0.40 clusters per µm^2^ in unstimulated cells to 0.18 clusters per µm^2^ in cells showing a speck (**Figure 2D**). This is consistent with the observed ASC recruitment (**Figure 2A**). Next, from the detected clusters, we investigated the number of localizations per cluster and the cluster size by calculating the radius of gyration (*R_g_*). We found the size of non-speck incorporated clusters to be very similar between unstimulated and stimulated cells (*R_g_* < 20 nm and localizations/cluster < 60) (**Figure 2E, F**). However, in speck-containing cells, an additional cytoplasmic cluster population appears with a larger size and more localizations (*R_g_* 20-80 nm and localizations/cluster > 60). We also applied a Ripley’s K clustering analysis ^49^ as an alternative approach for characterizing the spatial distribution of non-speck incorporated ASC. This analysis (**Figure 2G**) confirmed the decrease in cytoplasmic cluster density in cells containing a speck as shown by the shift of the amplitude of the obtained L(r)-r curve towards higher values and an increase of the L(r)-r maximum value consistent with a small population of larger clusters.

### Super-resolution microscopy reveals distinct morphologies of the endogenous ASC speck

Next, we investigated the nanoscale organization of the ASC speck itself ^50, 51^. We labelled endogenous ASC using a primary antibody in combination with a secondary F(ab’)_2_ fragment. A large proportion of the specks appeared as round, amorphous structures with a diameter of about 1 µm exhibiting a rough surface with short protrusions (**Supplementary Figure S4A**). Interestingly, for a large number of the specks, the higher resolution obtained by dSTORM imaging resolved ASC filaments reaching out from the dense core of the speck (**Figure 3A**). The number of clearly resolved filaments per structure varied but we rarely observed more than ∼10. We measured the diameter of the filaments using a Gaussian fitting of the intensity profile of multiple cross sections along each filament. The filament diameter was derived from the full width half maximum of the intensity profile, where we found a median value of 37.1 nm (**Figure 3B**). Considering the size of a primary/secondary F(ab’)_2_ antibody complex (∼11 nm) ^52^ used for labeling and the experimental localization precision of ∼ 10 nm (see Materials and Methods), the measured value corresponds to an actual thickness of the filament of ∼15 nm. When compared to values obtained for filaments formed by ASC *in vitro* and studied by EM (16 nm) ^17^, this value indicates that the majority of filaments are isolated single filaments.

**Figure 3.**
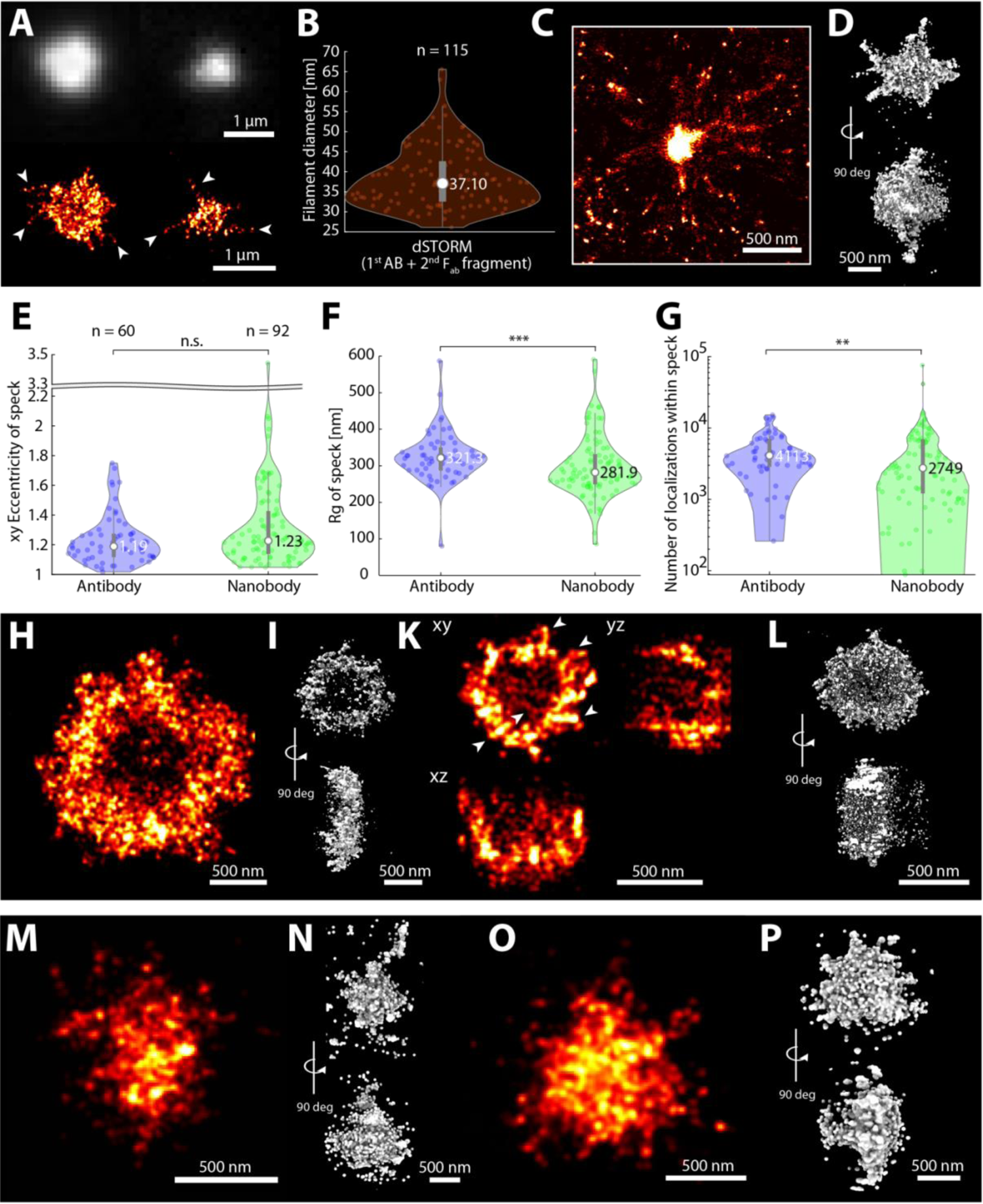
Super-resolution imaging of the endogenous ASC speck. A) Two ASC specks stained with primary antibody and secondary F(ab’)_2_ fragment in THP-1 cells exhibiting different sizes are shown after diffraction-limited widefield imaging (upper images) together with their 2D dSTORM reconstructions (lower images). B) The distribution of ASC filament diameters as measured in 2D dSTORM reconstructions on multiple cross-sections per filament. The circle in the violin plot indicates the median of the distribution. Data was obtained on three independent cell preparations and 18 individual specks. C) An ASC speck in BlaER1 cells imaged using DNA-PAINT super-resolution microscopy. Filaments are clearly observed reaching out from a dense core. D) A 3D dSTORM reconstruction of a single ASC speck stained with primary antibody and secondary F(ab’)_2_ fragment in THP-1 cells. E–G) Comparison of the endogenous ASC speck parameters determined using primary antibody and secondary F(ab’)_2_ fragment labeling (blue) and nanobody labeling (green). From the 2D dSTORM images, violin plots of the eccentricity (E), the radius of gyration (F) and the number of localizations (G) were determined. White circles in the violin plots indicate the median of the distribution. Statistical significance was assessed by a two-sided Two-Sample Kolmogorov-Smirnov Test (*) p<0.05, (**) p<0.01, (***) p<0.001. Data on antibody- and nanobody-stained specks were obtained from three and two independent cell preparations, respectively. H-L). A characteristic 2D reconstruction (H) and a 3D reconstruction of an ASC speck (I) as well as a z-stack projection of a 3D dSTORM reconstruction (K) and a 3D reconstruction of another ASC speck (L). Both exhibit a ring-like structure. In panel K, the arrows highlight filament-like structures. M-P) ASC specks stained with anti-ASC nanobodies. 2D (M) and 3D (N) reconstructions of two individual ASC specks imaged by dSTORM. O-P) 2D (O) and 3D (P) representations of an ASC speck imaged using DNA-PAINT. Short filamentous structures are resolvable at the edge of the dense speck core.

To validate these results, we also imaged the endogenous ASC specks using DNA-PAINT super-resolution microscopy ^53, 54^. Here, we used BlaER1 cells, transdifferentiated into monocytes/macrophages in which ASC was endogenously tagged with TagRFP ^55–57^. This alternative system again revealed the morphology of the endogenous ASC speck including filaments and a dense core. Strikingly, the filaments were much longer than the ones observed by dSTORM in THP-1 cells (**Figure 3C** and **Supplementary Figure S4B**).

Next, we performed 3D dSTORM imaging of the endogenous ASC specks. Consistent with the 2D dSTORM images, the specks had an overall spherical shape with filaments occasionally reaching out from a dense core (**Figure 3D, Supplementary Figure S5, Supplementary Figure S6, Supplementary Movie S1** and **Supplementary Movie S2**). The two-dimensional eccentricity for the ASC specks was determined to be 1.19, corresponding to more or less round specks as observed by eye, with a median radius of gyration of 321 ± 30.9 nm Median absolute deviation of the median (MAD) (**Figure 3E, F**, and Material and Methods). For some ASC specks, we observed a ring-like appearance (**Figure 3H - L** and **Supplementary Figure S4A + Supplementary Figure S6**). Some of the structures appeared less extended along the z-axis resulting in an overall disk-like shape (**Figure 3I + Supplementary Movie S3 + Supplementary Movie S4**) while others had a spherical, hollow shape (**Figure 3K + L, Supplementary Movie S5**). Although some specks appeared ring-like, the signal was not entirely excluded from the center. We hypothesized that local density differences may be responsible for the ring-like signal, perhaps due to the steric exclusion of primary and secondary antibody complexes from the dense center of the structure.

To test this hypothesis, we performed 3D super-resolution imaging using a 3-fold smaller ASC nanobody ^26^ to stain endogenous specks. Following nanobody labelling and dSTORM imaging, the specks appeared as amorphous structures with no obvious organization (**Figure 3M – P, Supplementary Figure S7** and **Supplementary Figure S8**). The structure remained spherical (**Figure 3E**) while the overall size of the specks, determined from the radius of gyration, was smaller (**Figure 3F**) and the total number of localizations within the speck decreased compared to specks stained with primary antibody and secondary F(ab’)_2_ fragment (**Figure 3G**). None of the 134 structures observed exhibited a hollow center. DNA-PAINT confirmed our observation of a smaller speck size and also resolved short filamentous extensions at the edge of the structure (**Figure 3O + P, Supplementary Figure S7B**, and **Supplementary Movie S6**). The absence of ring-like structures after nanobody labelling is consistent with our hypothesis that the dense regions within the speck limit the accessibility of the labeling probe. The smaller radius of gyration cannot be entirely explained by the high labeling density in the center, and we attribute it to the comparably low binding-affinity of the nanobody (apparent binding constant: 159.5 ± 1.5 nM ^26^). The nanobody has a single binding site, compared to two for a normal primary antibody, potentially explaining the higher labeling efficiency of the outer, lower-density regions of the speck by the antibody.

### Two-color super-resolution imaging confirms accessibility differences within endogenous ASC specks

To examine whether both the high-density core structure and low-density filaments are present on the same ASC speck, we performed two-color super-resolution microscopy using both nanobody and antibody labeling. Diffraction-limited widefield imaging showed that both labels specifically stained the ASC speck (**Figure 4A**). Consistent with our previous observation, the signal resulting from nanobody staining was smaller compared to the one obtained from antibody staining. Dual-color dSTORM reconstructions revealed that the antibody-labeled specks have a diameter of about one micrometer in widefield and an *R_g_* of 319.7 ± 29 nm MAD (compared to 324 ± 36 nm MAD under single-labeling conditions, **Table 1**) with the nanobody staining localized in the center of the speck (**Figure 4B** and **Supplementary Figure S9**). Antibody labeling also resolved filaments and, as observed in the single-color antibody staining, a subset of the ASC specks appeared hollow (**Figure 4C**). Strikingly, in these particles, the nanobody staining was more compact, with a significantly smaller *R_g_* (134 ± 50 nm MAD, compared to 282 ± 36.5 nm MAD under single labeling conditions, **Table 1**). We speculate that the higher-affinity antibody outcompetes the nanobody in the lower density regions of the speck, while only the nanobody is small enough to penetrate the dense core of the inflammasome. Aligning the dual-color structures along the center of mass of the antibody signals confirmed the observation that nanobody staining is confined to the center while the antibody complex was found more towards the speck periphery (**Figure 4D**). Hence, different labeling approaches bring out different features of the ASC speck. While it is typically beneficial to use the smallest available labels ^58, 59^, other factors such as binding affinity and density of the target structure can also play a role.

**Figure 4.**
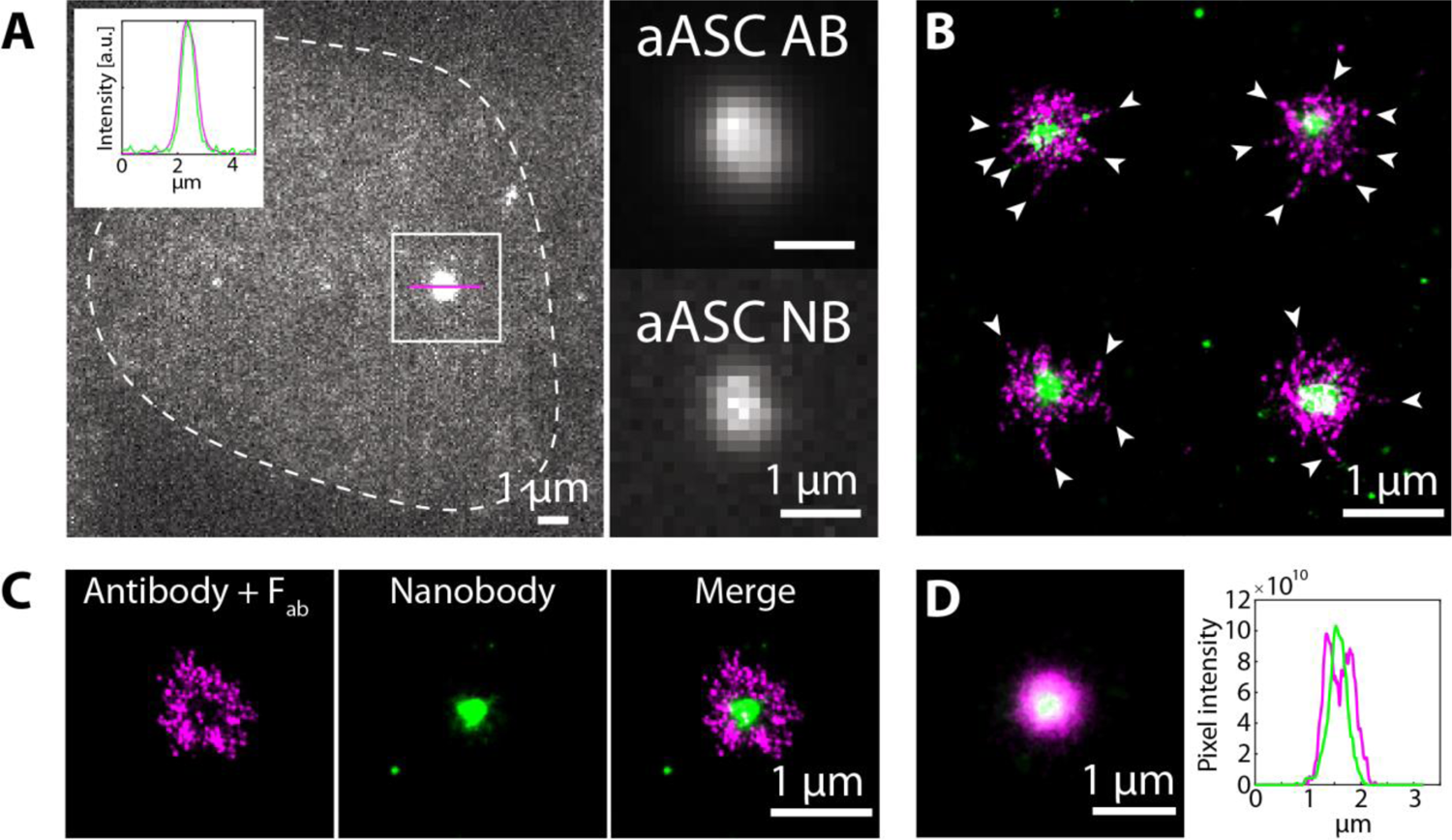
Dual-color dSTORM imaging of ASC specks simultaneously stained with primary antibodies/secondary F(ab’)_2_ fragments and nanobodies against ASC. A) Diffraction-limited widefield imaging of an ASC speck stained with primary antibody and Alexa Fluor 647-conjugated secondary F(ab’)_2_ fragment (the signal of which is shown here) and DyLight 755-conjugated nanobody. The dashed line shows the cell outline. The intensity profile of the ASC speck normalized to the peak of the intensity distribution along a cross-section of the speck (magenta line) is shown in the top left of the image (antibody signal: magenta; nanobody signal: green) showing that both staining approaches stain the same structure. The right part of the panel shows the boxed area split into the signal after antibody (top) and nanobody (bottom) staining. B) Four representative dual-color dSTORM reconstructions of ASC specks stained simultaneously with both antibodies and nanobodies (magenta: Alexa Fluor 647; green: DyLight 755). Arrows point towards filaments reaching out from the dense speck core. C) Dual-color dSTORM reconstruction of a speck showing a ring-like appearance after antibody + F(ab’)_2_-staining (left) labeling, staining of the dense core by the nanobody (center) and the merge of both labeling strategies (right). D) Alignment of dual-color-labeled specks (n = 35) along the center of mass of the antibody + F(ab’)_2_ fragment staining (left) and the intensity profile along a cross-section through the aligned structure (right). Data was obtained on a single cell preparation.

**Table 1.** Summary of measurement parameters on unstimulated cells and LPS + Nigericin-treated cells containing an ASC speck *All values occurred just once; MAD: Median absolute deviation of the median; AF 647: Alexa Fluor 647

|  |  | unstimulated cells | LPS + Nigericin-treated, Speck-containing cells |  |  |  |  |
| --- | --- | --- | --- | --- | --- | --- | --- |
|  |  | Primary antibody + secondary AF 647-conjugated F(ab')<sub>2</sub> fragment | Primary antibody + secondary AF 647-conjugated F(ab')<sub>2</sub> fragment w + w/o additional DL755-conjugated nanobody | Primary antibody + secondary AF 647-conjugated F(ab')<sub>2</sub> fragment only | Primary antibody + secondary AF 647-conjugated F(ab')<sub>2</sub> fragment w additional DL755-conjugated nanobody | AF 647-conjugated nanobody only | DL 755-conjugated nanobody w additional primary antibody + secondary AF 647-conjugated F(ab')<sub>2</sub> fragment staining |
| # of cells or specks |  | 7 | 57 (cluster analysis)<br>60 (speck analysis) | 21 | 39 | 92 | 42 |
| # of clusters |  | 2,514 | 15,566 | - | - | - | - |
| Mean cytoplasmic clusters/area [1/μm<sup>2</sup>] |  | 0.18 | 0.40 | - | - | - | - |
| Mean # of localizations per cluster |  | 23.19 | 30.08 | - | - | - | - |
| Median cluster size (2D-R<sub>g</sub>) [nm] |  | 14.86 | 20.43 | - | - | - | - |
| Mode cluster size (2D-R<sub>g</sub>) [nm] |  | 6.04 | 7.05 | - | - | - | - |
| xy Eccentricity of speck | Mean | - | 1.23 | 1.14 | 1.29 | 1.33 | 1.5 |
|  | Median | - | 1.19 | 1.13 | 1.24 | 1.23 | 1.4 |
|  | Mode | - | 1.02 | 1.02 | 1.04 | 1.07 | 1.03 |
|  | MAD | - | 0.08 | 0.04 | 0.07 | 0.11 | 0.19 |
| 2D-R<sub>g</sub> of speck [nm] | Mean | - | 325.2 | 332.6 | 321.2 | 298.1 | 158.3 |
|  | Median | - | 321.3 | 324.0 | 319.7 | 281.9 | 133.9 |
|  | Mode | - | 80.4 | 80.4 | 242.1 | 310.6 | 19.9 |
|  | MAD | - | 30.9 | 35.9 | 29.0 | 36.5 | 49.9 |
| Number of localizations per speck | Mean | - | 5010 | 7209 | 3826 | 5338 | 3848 |
|  | Median | - | 4113 | 7180 | 3307 | 2749 | 2787 |
|  | Mode | - | N/A* | 1062 | N/A* | 1297 | 345 |
|  | MAD | - | 1430 | 2691 | 954 | 1872 | 2061 |

### ASC forms a scaffold that increases in density with time

From the wealth of information that we gathered using widefield and super-resolution microscopy, we further quantitatively analyzed our data. Since the labeling of ASC with nanobodies was less robust, we limited our analysis to the data collected using primary antibody and secondary F(ab’)_2_ fragment labeling. We first manually segmented the cell and the speck so that we could calculate parameters dependent on the characteristics of the cells and the specks they contained. Widefield images were collected before super-resolution microscopy was performed. Hence, the information from both imaging modalities was available from the same cells. By plotting the total widefield intensity as a function of cell area, a clear correlation was observed (**Supplementary Figure S10A**) showing that larger cells express more ASC protein. Similarly, there is a positive correlation between the total number of localizations and cell size (**Supplementary Figure S10B**). In fact, the total number of localizations measured using dSTORM correlates well with the total widefield intensity, as one would expect (**Supplementary Figure S10C + D**). Small corrections for day-to-day variations were performed as discussed in the material and methods (**Materials and Methods** and **Supplementary Figure S11)**. In contrast, the intensity normalized by the cell area is relatively constant (**Supplementary Figure S10E**), suggesting that the concentration of ASC is constant across different cells.

Next, we investigated how the size of the speck varies with cell properties. The size of the speck, determined either by manual segmentation of the speck or via the calculation of the 2D radius of gyration, was found to increase with cell size (**Supplementary Figure S10F, Figure 5A**). To investigate how the radius of gyration depends on other parameters, we normalized out the cell-area dependence (**Supplementary Figure S10K** see **Materials and Methods**). Interestingly, the radius of gyration only weakly depends on the amount of ASC within the speck (**Supplementary Figure S10L, M**). The same trend was found when we quantified the speck size manually via the occupied area **(Supplementary Figure S10G - I)**.

**Figure 5.**
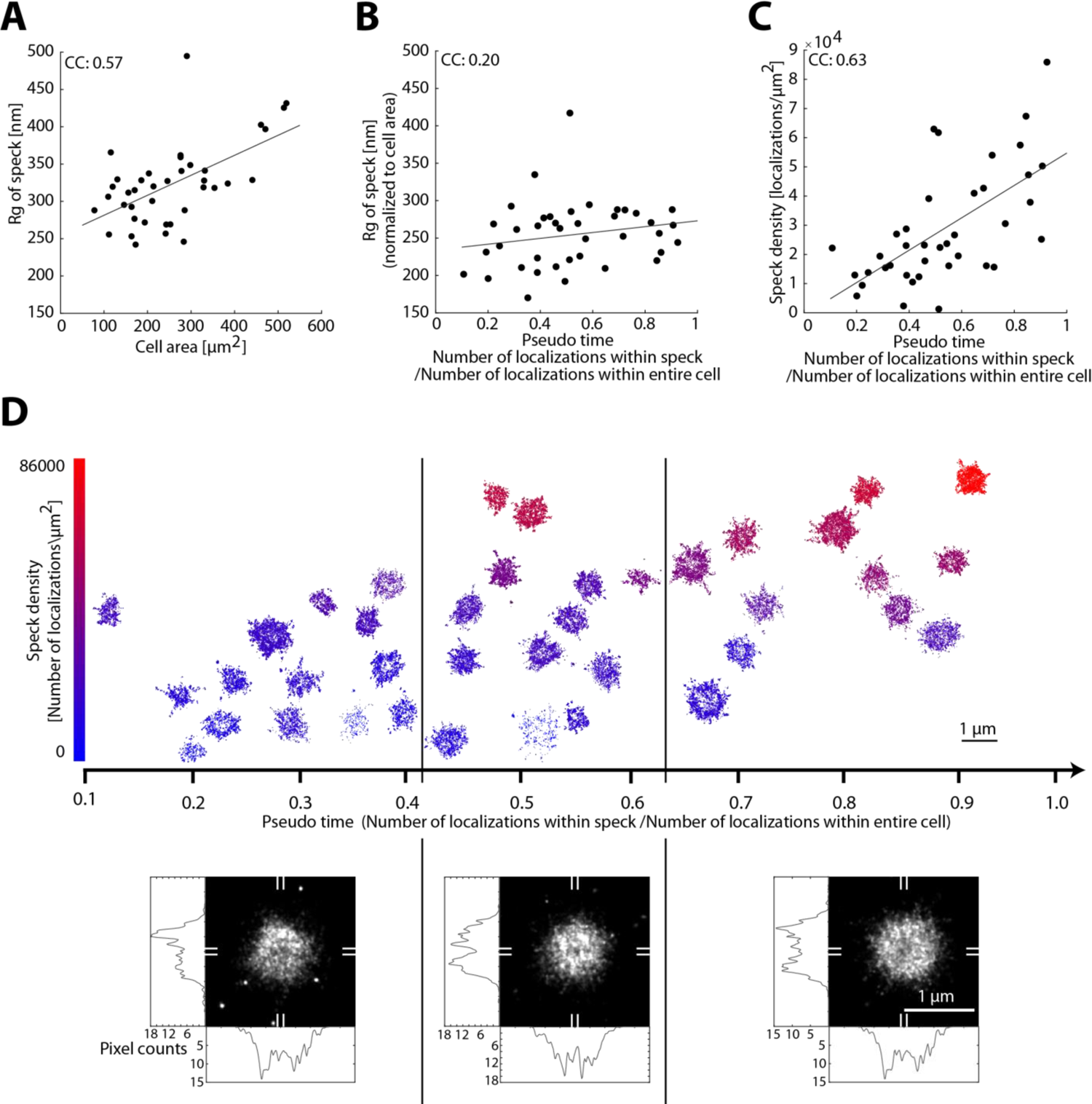
Dynamics of endogenous ASC speck formation. A) – C) Scatter plots of speck size (measured as the radius of gyration) as a function of the cell area (A), the speck size, normalized to cell area versus the fraction of the total localizations located within the speck (pseudo time) (B) and the speck density (measured as number of localizations per µm^2^) as a function of pseudo time (C). Larger cells form larger specks and the normalized speck size stays almost constant with increasing pseudo time whereas the speck density increases with pseudo time. CC: Pearson correlation coefficient. D) Plot of individual speck structures as a function of the pseudo time. The speck density is color-coded (scale bar is shown on the left) and specks are positioned in the vertical direction approximately according to their density. Specks were separated into three time bins (separated by the vertical black lines) and the super-resolution structures aligned and summed (lower panels). Horizontal and vertical cross-sections are shown (determined by averaging along the marked 9 pixel wide regions indicated on the periphery with white lines). The averaged speck structure appears more ring-like at late time points compared to early time points. Data was obtained on three independent cell preparations.

Speck assembly is a dynamic process, and recruitment of ASC to the perinuclear speck upon NLRP3 inflammasome activation occurs stochastically in the different cells we measured ^18, 46^. Thus, when fixed, different cells represent different stages of the assembly process. We used the ratio of ASC signal in the speck with respect to the total amount of ASC within the entire cell as a metric for the progression of speck assembly, i.e. pseudo time. For the ASC signal, we utilized the number of localizations as this is more reliable. We observed that the widefield intensity of the speck, as well as the number of localizations within the speck increased with ASC recruitment along the pseudo time axis, as expected (**Supplementary Figure S10N - O**). In addition, the cytosolic, non-speck ASC signal consistently decreased with the pseudo time regardless of whether we quantified it using the number of localizations, the cluster density or the localization density (**Supplementary Figure S10R - T**). A clear decrease in the integrated widefield intensity with the pseudo time is not observed (**Supplementary Figure S10Q**), we attribute this to the low level of signal remaining in the cytosol upon ASC recruitment to the speck, which suggests that the autofluorescence signal a significant fraction of the entire signal. A plot of the radius of gyration versus pseudo time showed little change in the size of the speck with pseudo time (**Supplementary Figure 10U**). This also holds true for the speck area (**Supplementary Figure 10V**), but the observation that the speck size depends on cell area could potentially confound the trend. Hence, we examined the normalized speck size as a function of pseudo time. The normalized radius of gyration of the speck (corrected for the correlation with cell area) increases only slightly during the course of ASC recruitment (**Figure 5B**). Similarly, when we normalized out the increase in speck area with cell size, only a small increase with pseudo time is observed **(Supplementary Figure S10P)**. We also plotted cell area with pseudo time (**Supplementary Figure S10W**), and observed no correlation. Although this was expected, it also verifies that the endogenous ASC speck formation is complete before pyroptosis and cell shrinkage is triggered. If the radius of gyration does not depend on pseudo time but the ASC content does, then we would expect the ASC density in the speck to increase with time. We investigated the speck density (localizations per area) and found it to be largely independent of the cell area (**Supplementary Figure S10X**) but clearly increasing with ASC recruitment (**Figure 5C**). In line with this observation, there is a clear correlation between the speck density and the amount of ASC within the speck, calculated either via the integrated widefield intensity within the speck (**Supplementary Figure S10Y**) or the number of localizations within the speck (**Supplementary Figure S10Z**). Hence, we conclude that the density and, to a much lesser extent, the size of the speck increases with ASC recruitment.

To visualize potential structural rearrangements of the speck, we plotted the endogenous ASC specks as a function of pseudo time (**Figure 5D**). No clear structural progression was visible. To look for more subtle changes during ASC recruitment, we divided specks into three groups according to the fraction of recruited ASC. We aligned the complexes in each group to their center of mass. The resulting sum projection showed an increased tendency to form a ring-like structure at later stages of recruitment, which is consistent with the observation of the specks become denser with time and thereby exclude the antibody from the center of the speck (**Figure 5D** lower panel).

## Discussion

The supramolecular NLRP3 inflammasome complex is a central component of the innate immune system, driving the maturation of the proinflammatory cytokines IL-1β and IL-18 as well as pyroptosis, a proinflammatory form of cell death. The protein ASC is critical during inflammasome formation and, upon cell stimulation, is recruited into a single, condensed structure, the ASC speck. ASC speck formation and organization are difficult to study due to the small size and heterogeneity of the complex. We used a combination treatment of LPS and nigericin, which led to robust induction of ASC specks with a broad distribution in ASC content (**Figure 1**). In diffraction limited wide-field microscopy, the complexes appear as spherical structures with very few morphological features. We then used dSTORM and DNA-PAINT, super-resolution techniques, to investigate the organization of endogenous ASC in the speck as well as in the cytoplasm. dSTORM measurements were performed on a total of 358 cells and 251 specks were analyzed in detail. This allowed us to reconcile several controversies regarding the reported structures of the ASC speck.

Our cluster analysis of the non-speck bound cytosolic ASC fraction revealed that the oligomeric size of the vast majority of ASC signals remained unchanged between unstimulated and speck-containing cells (**Figure 2E + F**). Considering the size of the labels, we measured an average *R_g_* in unstimulated cells of ∼15 nm (**Figure 2F**). This is in good agreement with an atomic force microscopy (AFM) study that measured the dimensions of full-length human ASC assemblies, which organized into disc-like oligomers of 1 nm in height and ∼ 12 nm in diameter *in vitro* ^11^. Upon stimulation, we observe a small increase in the number of larger clusters in the cytosol. This could be due to preassembly of a portion of ASC into higher-order ASC oligomers that later associate into the speck as has been suggested in the literature ^8, 60^. Western blot analysis of inflammasome activated cells found different levels of ASC multimerization in addition to the ASC speck with a large proportion of the protein being dimeric ^35^.

In our systematic measurements of ASC speck formation, the cells either showed a single speck or the ASC protein appeared homogenously distributed throughout the cell. We did not observe any concentration dependence of ASC as a function of distance from the speck (e.g. **Figure 2C**), consistent with previous observations ^46^ made in cells overexpressing ASC. This is understandable since the entire pool of ASC is recruited into the ASC speck within a few minutes during complex formation ^8, 18, 46, 61^. We were still able to detect cytosolic ASC in speck containing cells. The percentage of ASC recruitment we observed was typically 20% or above (with one exception) suggesting that the initial recruitment of ASC is faster than the fixation processes. This is consistent with live-cell imaging measurements in where the increase in speck size occurred over a time interval of ∼100 s ^46^. Our pseudo time observations indicate that it is mostly the speck density but not the speck size that increases with the percentage of ASC being recruited during the later stages of assembly.

The specks we recorded were smaller in size than those measured previously upon overexpression of ASC ^22, 25^. This is in line with our observation that the radius of gyration increases with total ASC content (**Figure 5A**). Moreover, NLRP3 inflammasome formation has recently been shown to occur at the microtubule organizing center (MTOC) ^62^ and a correlation has been observed between the size of the MTOC and the cell size (at least in *C. elegans*)^63^. This could provide an additional explanation for the size dependence of the ASC speck, in the case that the MTOC provides a scaffold for assembly of the speck. To reliably quantify the attributes of the amorphous, heterogeneous speck, we combined the results from 251 specks. The endogenous ASC speck has a size variation, determined from the 2D radius of gyration, ranging from 250 to 500 nm with an average radius of gyration of 321 ± 31 nm MAD (**Figure 3F**). The radius of gyration of the ASC speck corresponds the previously published value of ∼ 600 nm diameter for the endogenous structure (measured from cross-sections through the speck) ^35^.

Typically, one would expect the size of the speck to increase during assembly and hence the distribution of sizes to depend, in part, on the stage of assembly at which they are measured. However, when calculating the amount of ASC located within the speck relative to the total amount of ASC within the cell (i.e. our pseudo time), we found that the speck size depends only to a minor degree on the amount of recruited ASC (**Figure 5B**).

We found the majority of specks appeared as amorphous objects with an overall spherical structure, as determined from the calculated eccentricity. These results are consistent with previous observations ^8, 29^. In addition, we observed that some specks exhibit filamentous extensions protruding from the dense core. Although several studies suggested that the ASC speck is made up of intertwined filaments ^20, 22, 26, 27^, this has not yet been confirmed for the endogenous, unperturbed structure inside cells. We observe a dense core with filaments protruding from the endogenous ASC speck, which would be consistent with this hypothesis.

Although we could not analyze the observed filaments in detail due to the limited labeling density, we did observed filaments of varying length and thickness. Some specks exhibit many short and faint fibrils (**Figure 3 M-P**) while others show a small number of longer filaments protruding from the edge of the speck center (**Figure 3C**). The thin fibrils are reminiscent of the fibrils resolved by EM at the edge of the *in vitro* formed ASC assembly ^22^. A similar variation in thickness of the filaments protruding from the dense speck core has been observed in immortalized ASC^-/-^ bone marrow-derived macrophages (BMDMs) ^20^. Interestingly, electron microscopy data of *in vitro* assembled structure from full-length human ASC protein suggested that individual ASC filaments can laterally stack via the exposed CARD domains and the authors raise the question of whether this could also happen in the endogenous structure ^17^. The average filament thickness we obtained on the above-mentioned filaments agrees with the thickness reported for a single filament (∼16 nm) ^17^ suggesting that, for the majority of filaments, lateral stacking does not take place.

It is important to mention that the majority of our experiments were conducted in caspase-1 knock-out cells. Hence, we cannot exclude that the lack of caspase-1 binding to ASC has an influence on the appearance of the speck. Experiments performed using DNA-PAINT on BlaER1 cells that express caspase-1 showed the same dense, amorphous core with protruding filaments. This strongly suggests that the measured structures are indicative of the morphology of the ASC speck and is consistent with what has been observed in experiments in cells with ASC overexpression ^20, 22, 25^. However, in comparison to the previous observations of the speck formed after overexpression ^22, 25^, we found fewer and shorter filaments protruding from the endogenous ASC speck core with the majority of filaments being 500 nm long or shorter. It is conceivable that overexpression leads a larger number of filaments emanating from the formed speck. It is also possible that caspase-1 influences the filament length as our measurements in BlaER1 cells that express caspase-1 showed longer filaments than the ones observed in THP-1 caspase-1 knock-out cells (**Figure 3C** and **Supplementary Figure S4B**). The longer filament lengths are in line with a previous report where *in vitro* experiments measured filament lengths of 500-2000 nm for full length mouse ASC ^16^. However, we cannot entirely rule out that the differences in filament length are due the details of the super-resolution approach used.

An overall ring-like appearance of the ASC speck has been proposed since its discovery ^3^. However, the question of how the ring-like assembly and the irregular structure made up of intertwined filaments relate to each other remains unanswered ^39–42^. Here, we provide new data that offers an explanation for the observed ring-like structure. Complementing previous studies, which showed a ring-like assembly of ASC following antibody labelling, we used labels of different sizes, primary antibodies labeled with secondary F(ab’)_2_ fragments and nanobodies. While primary antibody plus secondary F(ab’)_2_ fragment labeling occasionally showed ring-like structures, simultaneous labeling with nanobodies revealed a dense core in the speck (**Figure 4**). Hence, we conclude that the endogenous speck can be divided into two different regions: a dense core and a less dense periphery (**Figure 6**). This would imply that the ring shape originates from the primary and/or secondary antibody being less likely to penetrate into the dense center of the structure and thus leading to a decrease in labeling efficiency at the center of the structure. Conversely, the less dense structures at the periphery are less efficiently labeled by the nanobody as it has only one binding site per protein with, in this case, a relatively low binding affinity (∼160 nM). This also demonstrates the importance of verifying super-resolution structures using different labeling strategies and approaches.

**Figure 6.**
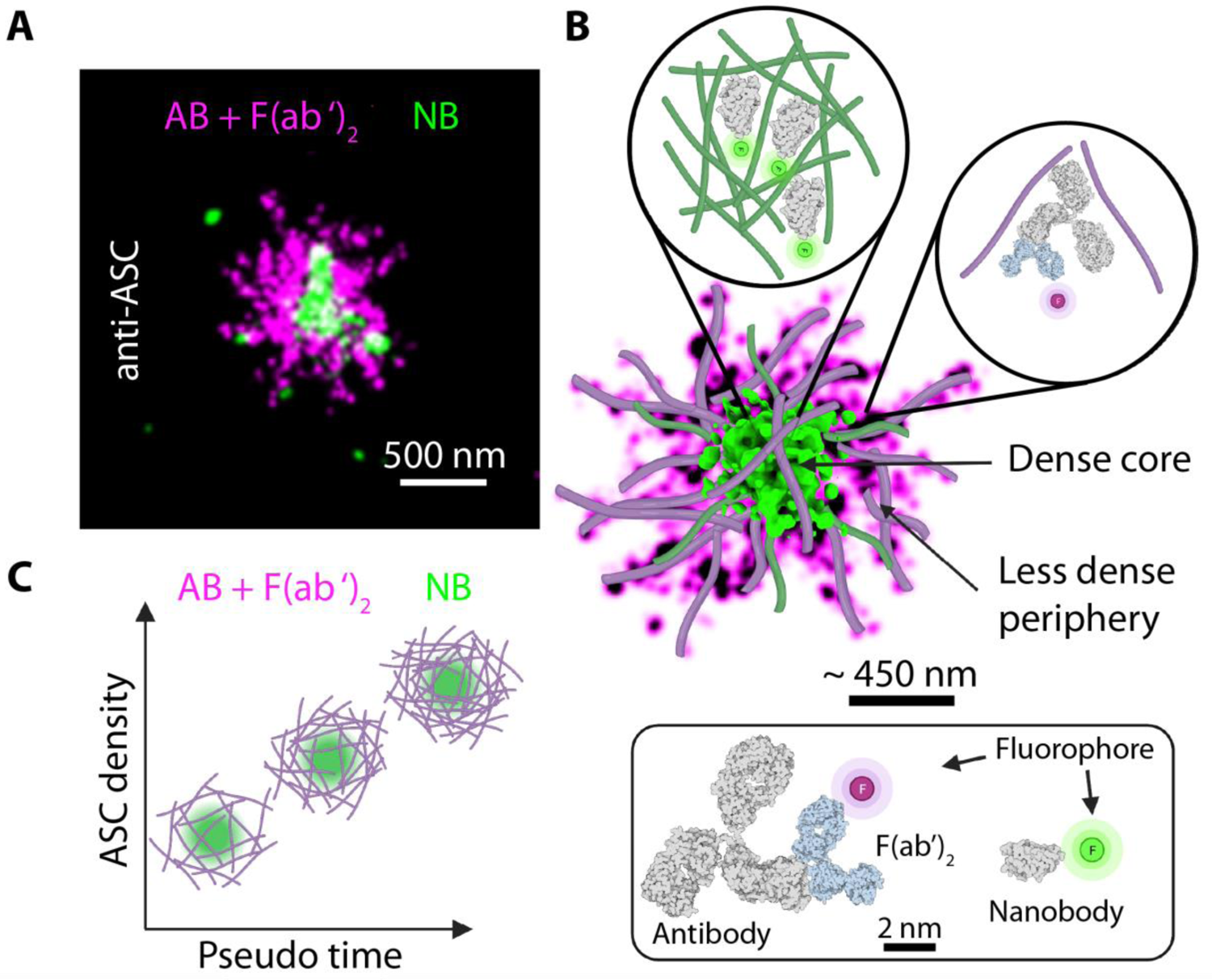
Model of endogenous ASC speck formation. A) 2-color dSTORM reconstruction of an ASC speck stained with primary anti-ASC antibody and secondary Alexa Fluor 647-conjugated F(ab’)_2_ fragment (magenta) and a DyLight 755-conjugated anti-ASC nanobody (green). The nanobody signal is concentrated in the center of the structure whereas the primary antibody / secondary F(ab’)_2_ fragment staining is also observed in the periphery of the structure. B) Schematic model for the supramolecular structure of the ASC speck based on measured data overlaid with modeled filaments. The structure is characterized by a dense core and a less dense periphery. The zoom ins illustrate that the nanobody is able to penetrate into the dense core of the speck whereas the antibody preferably labels the less dense periphery of the structure. The insert at the bottom of panel B illustrates the size difference between the labeling probes. C) Illustration of speck formation over time. The ASC speck forms from intertwined filaments assembling into a scaffold whose size scales with the cell size. Early stages of speck formation are characterized by a loose assembly into which the antibody (magenta) and nanobody (green) can penetrate. Further ASC recruitment into the speck leads to denser structures but only a marginal increase in its size. Antibody and F(ab’)_2_ fragment staining is sterically excluded from the dense center of the structure resulting in an overall ring-like appearance. The smaller nanobody stains the dense core of the speck but is washed away from the less dense regions of the structure due to its lower binding affinity. AB: Antibody; F(ab’)_2_: F(ab’)_2_ fragment; NB; Nanobody

## Conclusion

Taken together, we investigated the nanoscale organization of the endogenous ASC speck using super-resolution microscopy. We found that the speck size was heterogenous and correlated with cell size. The speck contained a dense core with filaments protruding from the center of the speck. This was observed for dSTORM and DNA-PAINT measurements in THP-1 cells as well as DNA-PAINT measurements in BlaER1 cells. These results are consistent with the model of the ASC speck being an assembly of intertwined filaments. By using differently sized labels, we found that the ring-like appearance of the speck is a result of the labeling approach and limited access of the primary antibody and/or F(ab’)_2_ fragment to the dense core of the speck. Conversely, the small nanobody was able to label the center of the speck but was less efficient in labeling the low density periphery regions due to the lower affinity of this nanobody (**Figure 4**). Hence, it is important to verify super-resolution structures using different labeling approaches. Finally, by measuring the fraction of recruited ASC, we hypothesize that speck formation starts with a loose scaffold that becomes denser but only marginally larger during ASC speck formation.

## Materials and Methods

If not stated differently all reagents were purchased from Thermo Fisher Scientific, Massachusetts, USA.

### Cultivation of cells and activation of the NLRP3 inflammasome

For the experiments performed with THP-1 caspase-1 knock-out cells, the THP-1 cells were cultivated at 37°C, 5% CO_2_ in Roswell Park Memorial Institute 1640 medium (21875034) supplemented with 10% (v/v) heat-inactivated fetal bovine serum (FBS) (10500064), 1 mM sodium pyruvate (11360039) and 100 U/ml Penicillin/Streptomycin (15140122), and maintained at a density between 1×10^5^ and 1×10^6^ cells per milliliter. One day prior to seeding the cells, coverslips (1.5, Menzel Gläser, 18 mm) were coated with 0.01% poly-l-ornithine solution (A-004-C, Merck-Millipore, Massachusetts, USA) in the dark. Cells were seeded at a density of ∼ 75×10^3^/cm^2^ in culture medium supplemented with 100 ng/ml Phorbol 12-myristate 13-acetate (PMA) (BML-PE160-0001, Enzo Life Sciences, Lörrach, Germany) and differentiated into macrophage-like cells for three days. To increase NLRP3 protein expression, cells were primed for three hours with 1 µg/ml Lipopolysaccharide (LPS) from *E. coli* K12 cells (Ultrapure tlrl-peklps, Invivogen, San Diego, California, USA). To suppress cell death due to caspase activity, cells were subsequently treated with 20 µM of the pan-caspase inhibitor Z-VAD (tlrl-vad, Invivogen) for 60 mins. The NLRP3 inflammasome response was activated by incubating the cells with 10 µM Nigericin (N7143-5MG, Sigma Aldrich, Missouri, USA) for 90 mins.

### ASC immunofluorescence with primary antibody and secondary F(ab’)_2_ fragment and for double-labeling with an ASC nanobody

All steps were performed at room temperature if not stated differently. Cells were washed once with phosphate-buffered saline (PBS) solution to remove remaining serum proteins, fixed for 15 mins in the dark with 4% paraformaldehyde (PFA) (E15710-S, Electron Microscopy Sciences, Pennsylvania, USA) in PBS. Subsequently, PFA was rinsed once with PBS and then quenched by rinsing once with 0.1 M NH_4_Cl (254134, Sigma-Aldrich) in PBS followed by a 15 min incubation with 0.1 M NH_4_Cl in PBS. Permeabilization and blocking was done for 30 mins in 10% normal goat serum (NGS) (16201), 0.5% Triton X-100 (T8787, Sigma-Aldrich) in PBS followed by a washing step with PBS and 30 mins incubation in Image-iTFX Signal enhancer (I36933). After two additional PBS washing steps, cells were incubated overnight at 4°C with purified monoclonal mouse anti-human ASC antibodies (TMS-1) (clone HASC-71) (Biolegend, California, USA) at a final concentration of 10 µg/ml diluted in 10% NGS + 0.5% Triton X-100 in PBS. 25 µl of this solution was applied to the fixed cells on a coverslip. The plasmid encoding for the ASC nanobody was provided by Prof. Dario Alessi, University of Dundee via the MRC – Protein Phosphorylation and Ubiquitylation Unit [DU 54832]. The nanobody encoding sequence with a C-terminal cysteine was amplified for labeling and expressed at the protein production core facility of the Max-Planck-Institute for Biochemistry in Martinsried, Germany. Parts of the nanobody were conjugated to Alexa Fluor 647 in house as described in ^64^ and at Nanotag Biotechnologies GmbH, Göttingen, Germany. In the case of dual-color labeling, DyLight755-conjugated anti-ASC nanobody was purchased from Nanotag and mixed into the antibody solution at 1 µg/ml final concentration. Non-specific sticking of antibody and nanobody was minimized by washing three times with 0.1% Triton X-100 in PBS. Labeling with secondary F(ab’)_2_-goat anti-mouse IgG (H+L) cross-adsorbed Alexa Fluor 647-conjugated F(ab’)_2_ fragment (A-21237) was performed at 200 ng/ml final concentration in 10% NGS + 0.5% Triton X-100 in PBS for 1 hr followed by three washing steps with 0.1% Triton X-100 in PBS and postfixation with 3% PFA in PBS for 10 mins. The specificity of the staining procedure was confirmed by staining THP-1 caspase-1 knock-out cells only with the F(ab’)_2_ fragment without previous administration of the primary antibody as well as by applying the staining protocol to THP-1 ASC knock-out cells (**Supplementary Figure S10**).

### ASC immunofluorescence with ASC nanobody

THP-1 caspase-1 knock-out cells were activated for the NLRP3 inflammasome, fixed and quenched as described above followed by permeabilization with 0.05% Saponin (47036, Sigma-Aldrich), 1% BSA (A7030, Sigma-Aldrich), and 0.05% NaN_3_ (S2002, Sigma-Aldrich) in PBS for 20 mins. Afterwards, the sample was wash for 2 mins with PBS and then the sample was blocked for 30 mins with Image-iTFX Signal Enhancer (R37107). The sample was then washed twice for 2 mins with PBS and stained against ASC with 1 ug/ml Alexa Fluor 647-conjugated nanobody and postfixed for 10 mins in 3% PFA in PBS. The specificity of the staining was confirmed using THP-1 ASC knock-out cells (**Supplementary Figure S13**).

### Confocal imaging

For the confocal imaging shown in Figure 1, the samples were embedded in ProLong Gold (P10144) on standard glass slides for at least one day before imaging. Microscopy was then performed on a Leica SP8 STED 3x equipped with a 470-670nm white light laser and a 100 x PlanApo /NA 1.4 objective. The spinning disk confocal microscopy images shown in **Supplementary Figure S12A** were measured on a Zeiss Cell Observer SD. Alexa Fluor 647 and DAPI were excited using a 639 nm and 405 nm laser, respectively. Fluorescence was separated using a 660 nm longpass filter and recorded on the two Evolve 512 electron-multiplying charge-coupled device cameras (Photometrics) of the system equipped with a 525/50 and a 690/50 bandpass filter, respectively.

### dSTORM imaging and analysis

Samples were imaged on a flat field-optimized widefield setup as described before ^51^. Briefly, lasers of 405 nm, 642 nm and 750 nm were expanded and reflected into a 60 x objective (CFI60 PlanApo Lambda 60x / NA 1.4, Nikon) using a custom appropriate dichroic mirror (ZT405/561/642/750/850rpc, Chroma). Fluorescence emission was imaged onto a sCMOS camera (Prime, Photometrics) using one of two emission filters (ET700/75M and ET810/90m, Chroma) combined with a short-pass filter (FF01-842/SP, Semrock) and a tube lens of 200 mm focal length. The microscope was controlled using Micromanager ^65, 66^. Widefield images were collected prior to dSTORM recordings. We typically recorded between 20-80k frames at 10 ms exposure time. For 3D dSTORM, we introduced a cylindrical lens (f=1000 mm, Thorlabs LJ1516RM-A) into the emission path. Single- and dual-color single-molecule localization microscopy (SMLM) imaging was carried out with an optimized SMLM buffer, as described previously ^67^. Single molecules were localized using an sCMOS-specific localizer routine introduced by Huang *et al.* ^68^ and included in a custom MatLab program used for data analysis. To exclude non-specific localizations, filter parameters were adjusted using datasets acquired using ASC knock-out THP-1 cells. Drift correction was performed using the Redundancy Cross-Correlation (RCC) algorithm introduced by Wang *et al.* ^69^. The cells and specks were manually segmented. Cytoplasmic localizations were clustered using DBSCAN where a cluster was defined as a group of 10 – 300 localizations within a search radius of 70 nm ^48^ or Ripley’s K function ^49^. Super-resolution images were reconstructed using ThunderStorm ^70^ with 10x magnification and applying a Gaussian blur of one pixel (10.6 nm). For dual-color data sets, the localizations in the DyLight 755 channel were registered to the ones in the Alexa Fluor 647 channel by an affine transformation calculated from widefield images of immobilized TetraSpeck Microspheres (T7279) on a plasma cleaned coverslip recorded in both channels. 3D representations were rendered using Chimera X ^71^.

The eccentricity, the radius of gyration and the number of localizations of the specks were calculated from the manually segmented localizations within the structure. The radius of gyration was calculated as the square root of the sum of the variances of the x and the y coordinates of the individual localizations within each speck. As, in most cases, the specks are amorphous, spherical structures and we have significantly better resolution in the radial dimension, we limited the calculated radius of gyration to two-dimensions. The eccentricity of the individual specks was determined from the localizations by first calculating the covariance matrix of the individual localizations. From the covariance matrix, the eigenvectors were calculated, which provide a measure for how elliptical the distribution of localizations is and calculates the direction of the major and minor axes of the ellipse. We then take the square root of eigenvectors, which give a measure of length for the different axes. Finally, the eccentricity is calculated by taking the ratio of the major axis (square root of the maximum eigenvector) to the minor axis (square root of the minimum eigenvector). Circular objects have an eccentricity near 1. For our microscope we determined an experimental localization precision of σ_xy_ = 12 nm for Alexa Fluor 647 and σ_xy_ = 21 nm for DyLight755 ^51^.

For the quantitative image analysis, measurements of the integrated widefield intensity were corrected by subtracting background counts determined individually for each field of view by averaging pixel counts in an area absent of cellular signal. To calibrate for the experimental differences observed between measurement days, the widefield intensity per cell area and the number of detected localizations per widefield intensity were corrected separately for each measurement day. For this, we used the correlation of fluorescence intensity with cell size and the number of localizations versus widefield intensity. The correlation was determined for each day by fitting the data with a line and the slope was scaled to correspond to the maximum of all experiment days. A correction factor was determined for each measurement day and used to rescale the measurements to the measurement day with the largest slope. The scaled widefield intensity and number of localizations were then used for the ensuing analyses. **Supplementary Figure S11** illustrates the applied correction procedure.

The cell area dependence of the radius of gyration (*R_g_*) was normalized out by first plotting the *R_g_* against the cell area and fitting the data with a line. Subsequently for each data point, the product of the slope of the line and the value of the cell area was calculated and subtracted from the *R_g_* value. The speck area was normalized analogously.

### BlaER1 cell preparation and DNA-PAINT imaging

BlaER1 ASC-TagRFP cells were cultivated as described for the THP-1 cells. Cells were transdifferentiation for 7 days at 60,000 cells/cm^2^ in medium containing 50 ng/ml IL-3 (200-03 B, Peprotech, New Jersey, USA), 50 ng/ml M-CSF (300-25 B, Peprotech) and 500 nM β-Estradiol (E8875, Sigma-Aldrich) in cell culture-treated plastic bottom slides (80826, ibidi, Gräfelfing, Germany). Cells were then trypsinized, transferred to glass bottom slides coated with poly-l-ornithine as described for THP-1 cells in cultivation medium without growth factors at the same density and incubated overnight at 37°C, 5% CO_2_. The next day, the NLRP3 inflammasome was activated by priming the cells for 14 hrs with 200 ng/ml Lipopolysaccharide (LPS) from *E. coli* K12 cells in medium followed by caspase inhibition with 20 µM Z-VAD-FMK incubation for 60 mins as described for THP-1 cells and 3 hrs incubation with 6.5 µM Nigericin. Cells were fixed and stained as described for THP-1 cells with the exception that a secondary donkey anti-mouse antibody (715-005-151, Jackson ImmunoResearch) conjugated to a P3 docking strand (TTCTTCATTA) was used instead of the F(ab’)_2_ fragment. Specks were identified by excitation of TagRFP with a 561 nm laser, followed by bleaching the signal and DNA-PAINT data were recorded using a P3-Cy3B imager strand (AATGAAGA-Cy3B) at 1 nM concentration and HiLo illumination at 1.2 kW/cm² excitation with the same laser at 100 ms camera exposure time for 8000 frames.

THP-1 cells imaged by DNA-PAINT were first labeled with a low amount of Alexa Fluor 647-conjugated anti-ASC nanobody to facilitate identification of the speck. Subsequently, the nanobody conjugated to a P3 docking strand was applied and the structure was imaged using the complementary imager strand labeled with Cy3B at 0.5 nM concentration. Images were recorded for 20,000 – 30,000 frames with an exposure time of 100 ms and a laser power density of ∼200 kW/cm². Super-resolution images were reconstructed using the Picasso [51] and ThunderStorm [64] softwares and rendered with a Gaussian blur of one pixel corresponding to 13 nm.

## Authors contributions

I. M. G. and D. C. L. designed experiments, I. M. G., C. S. and D. C. L. analyzed the data, I. M. G., G. A. and G. M. P. conjugated the nanobody, I. M. G. and G. M. P. prepared immunofluorescence samples, I. C. S. and I. M. G. recorded dSTORM data, S. S. and I. M. G. recorded DNA-PAINT data, C. S. prepared THP-1 knock-out cells, T. S. E. and C. S. helped with establishing the immunofluorescence protocols, II. D. C. L., V. H., S. M., R. J. and I. M. G. acquired funding, I. M. G., C. S. and D. C. L. wrote the manuscript with input from all other authors.

## Acknowledgements

We thank Fionan O’Duill for advice on cell culture and inflammasome activation. We would also like to thank Dr. Steffen Frey and Dr. Hansjörg Götzke from NanoTag Biotechnologies GmbH, Göttingen, Germany as well as Dr. Sabine Suppmann and Leopold Ulrich from the Protein Production Core Facility at the Max-Planck-Institute for Biochemistry, Martinsried, Germany for expressing and labeling nanobodies used in this study, and for fruitful discussions.

## Supplementary Information

**Supplementary Figure S1:**
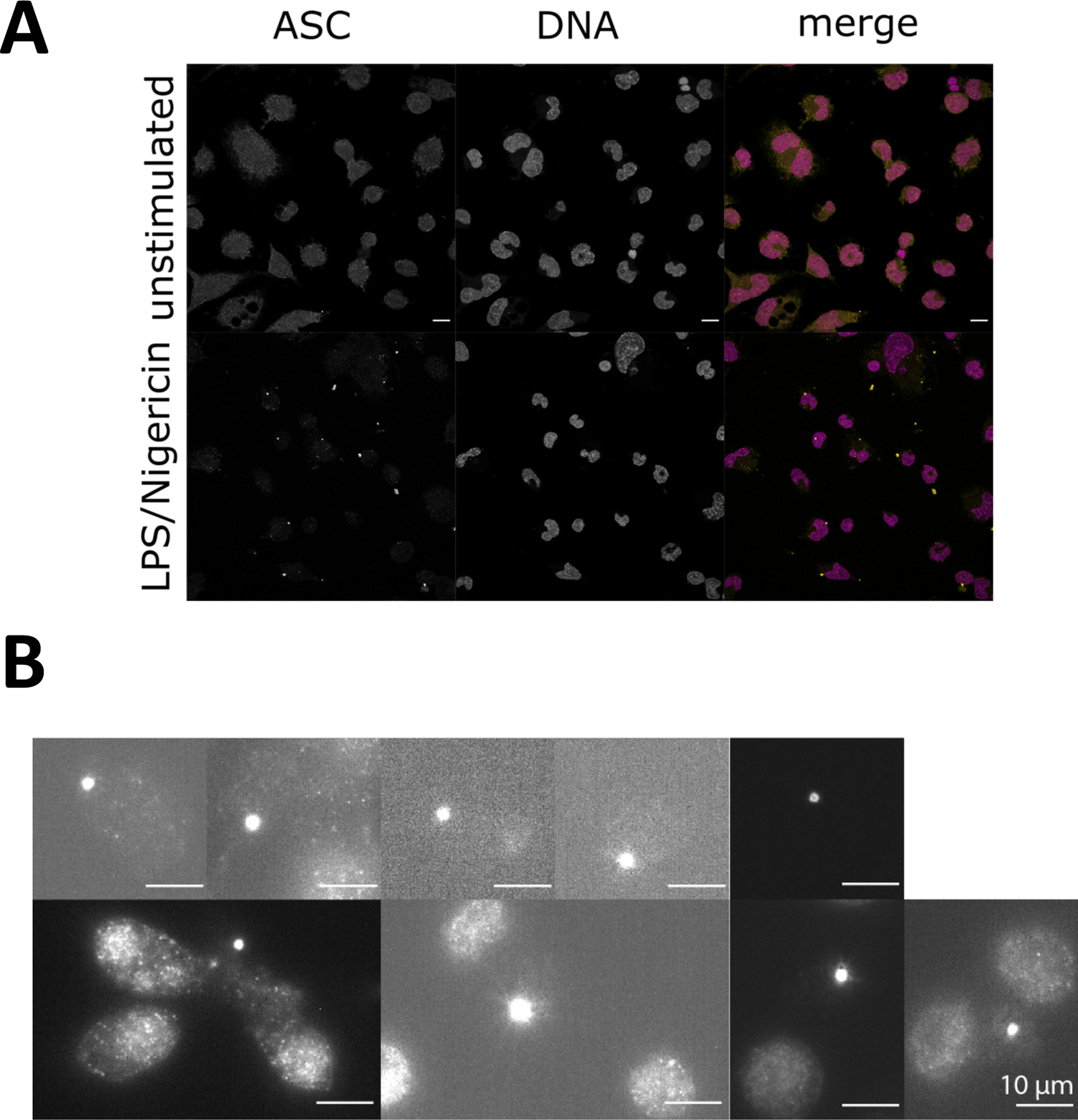
Diffraction-limited images of fluorescently labeled ASC in unstimulated and stimulated THP-1 cells and of extracellular ASC specks. A) Confocal images (maximum projections) of fluorescently labeled ASC in unstimulated (upper panel) and in LPS + Nigericin-stimulated THP-1 cells (lower panel). Images as shown in Figure 1. The scale bar corresponds to 10 µm. B) Widefield images of extracellular ASC specks stained with primary ASC antibody and secondary Alexa Fluor 647-conjugated F(ab’)_2_ fragment.

**Supplementary Figure S2:**
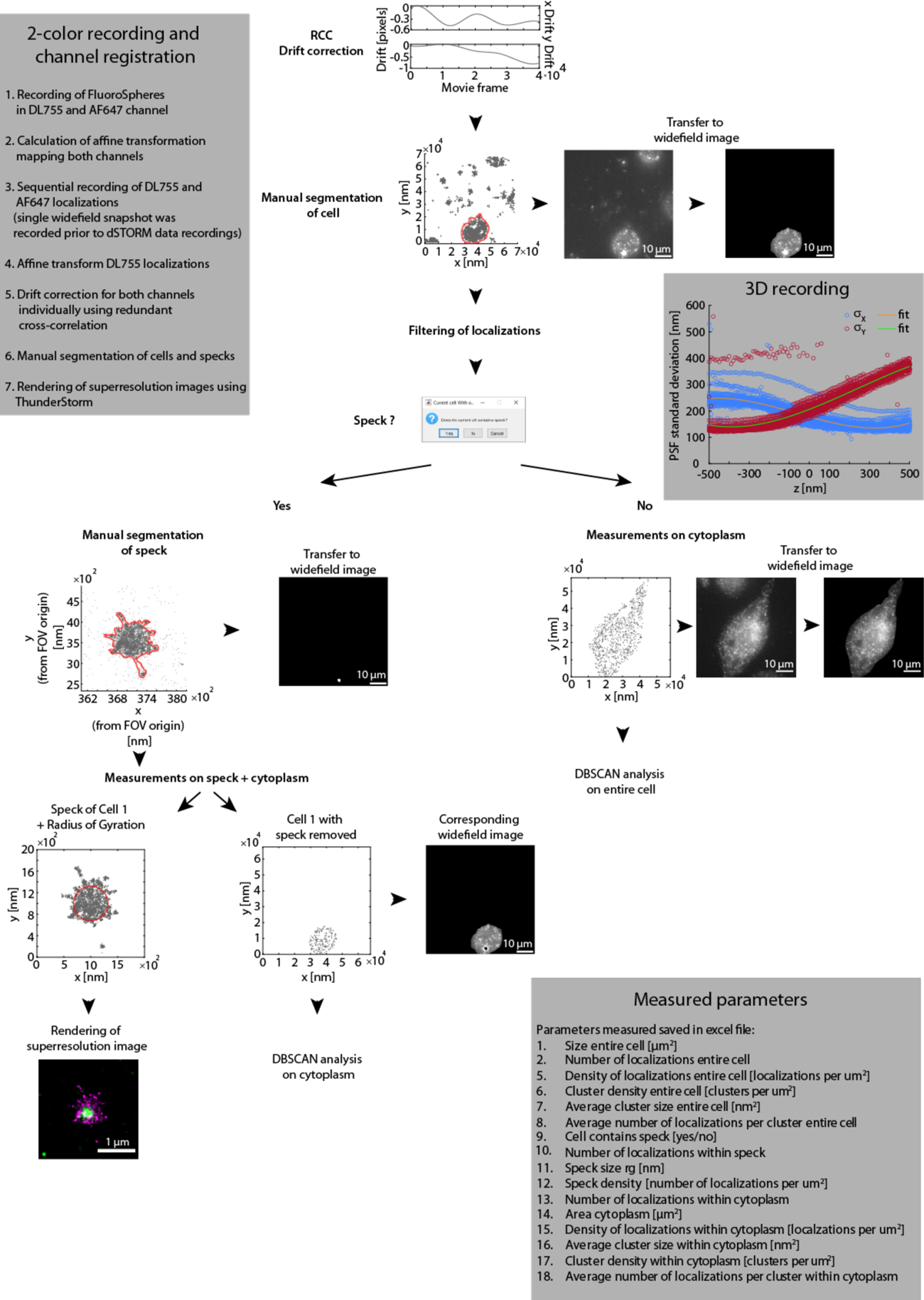
Applied image data analysis workflow. Widefield and dSTORM Microscopy experiments were performed on fixed cells. After localizing individual molecules and performing drift correction, the cells were manually segmented from the surroundings using the localizations and the obtained mask was applied to the widefield image. The localizations were filtered to remove unspecific and low quality localizations. In cells containing an ASC speck, the speck was manually segmented and the obtained mask was applied to the widefield image. A super-resolution image was rendered from the localizations. Finally, various parameters for the entire cell, the cytosol and the ASC speck were calculated (listed in the box in lower, right corner). Cytosolic localizations were analyzed using the DBSCAN analysis (Supplementary Figure S3). The axial positions for 3D images were obtained by recording and fitting a calibration curve measured on TetraSpeck Microspheres (see gray box to the upper right). The 2-channel registration was performed as described in the box in the upper left. RCC: Redundancy cross-correlation.

**Supplementary Figure S3:**
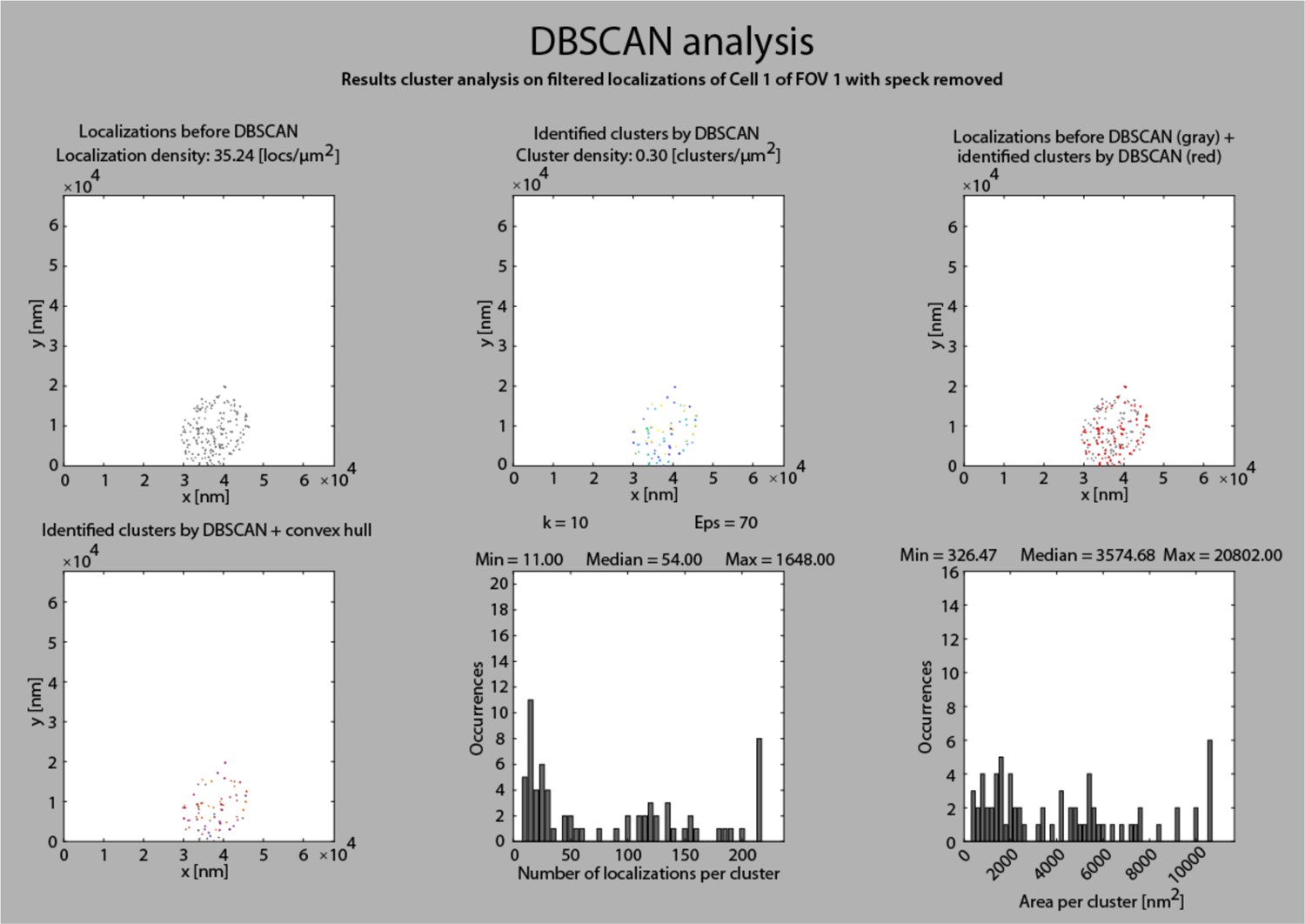
The DBSCAN analysis. A screen shot of the DBSCAN analysis of the cytosolic non-speck bound ASC signal of the cell shown in Supplementary Figure S2 is depicted. The localization density before DBSCAN is shown in the top left panel. The top middle panel depicts the identified clusters after an analysis using a search radius of 70 nm and 10-300 localizations per cluster. Individual clusters are color-coded. The top right panel depicts an overlay of the identified clusters (red) with the unclustered localizations (gray). The bottom left panel depicts the clusters together with their convex hull. The bottom middle and right panels show histograms of the number of localizations per cluster and the area per cluster derived by calculation of the convex hull, respectively.

**Supplementary Figure S4:**
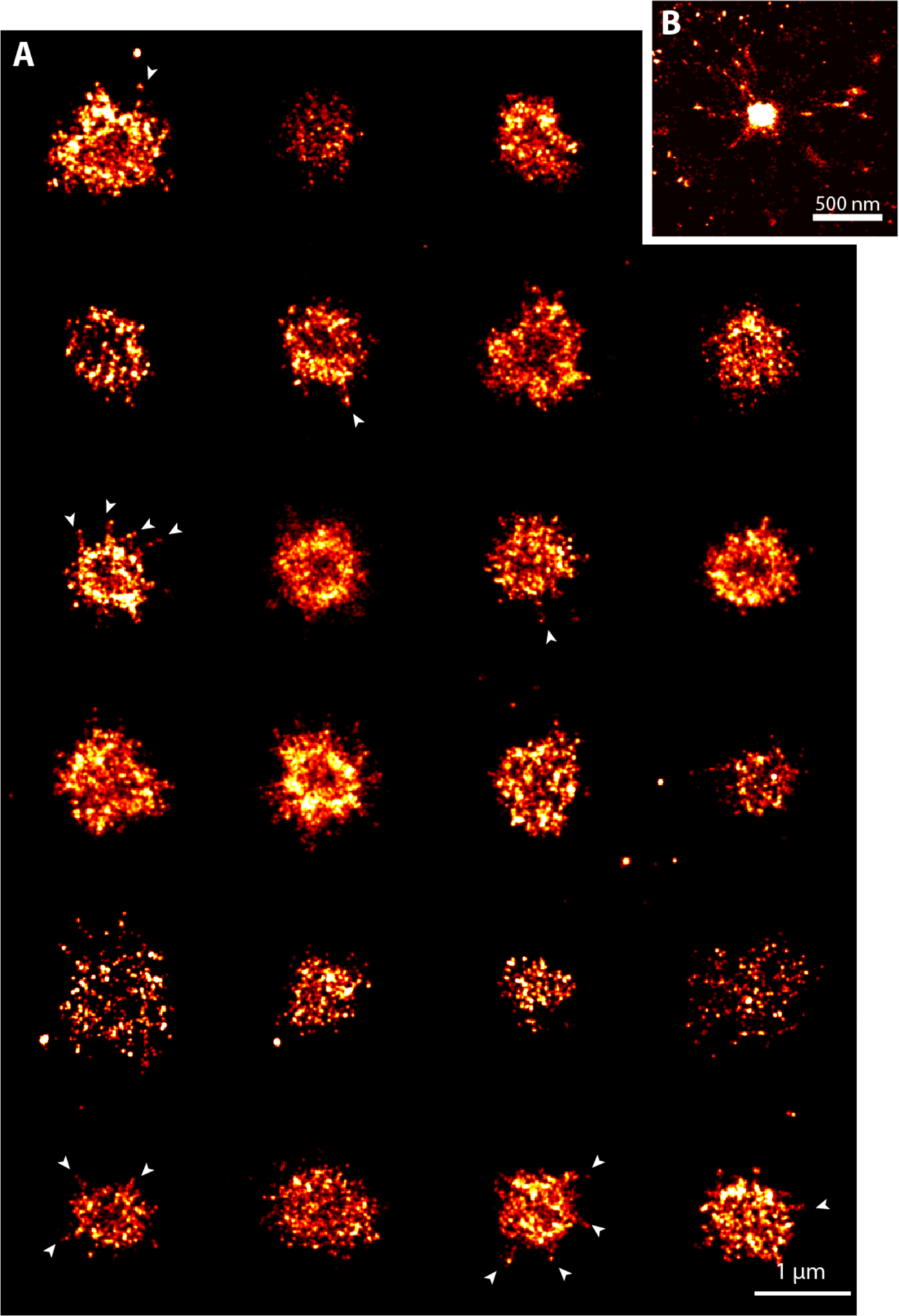
Two-dimensional super-resolution images of endogenous ASC specks. A) A gallery of endogenous ASC specks labeled with an ASC antibody and an Alexa Fluor 647-conjugated F(ab’)_2_ fragment in THP-1 cells imaged using dSTORM. Some structures appear as ring-like structures and other as dense amorphous complexes. In several structures, filamentous extensions are observed, which are indicated by the arrows. B) An ASC speck observed using DNA-PAINT labeled with an ASC antibody and a secondary antibody conjugated with a P3 docking strand for DNA-PAINT imaging. Data was obtained on three independent cell preparations.

**Supplementary Figure S5:**
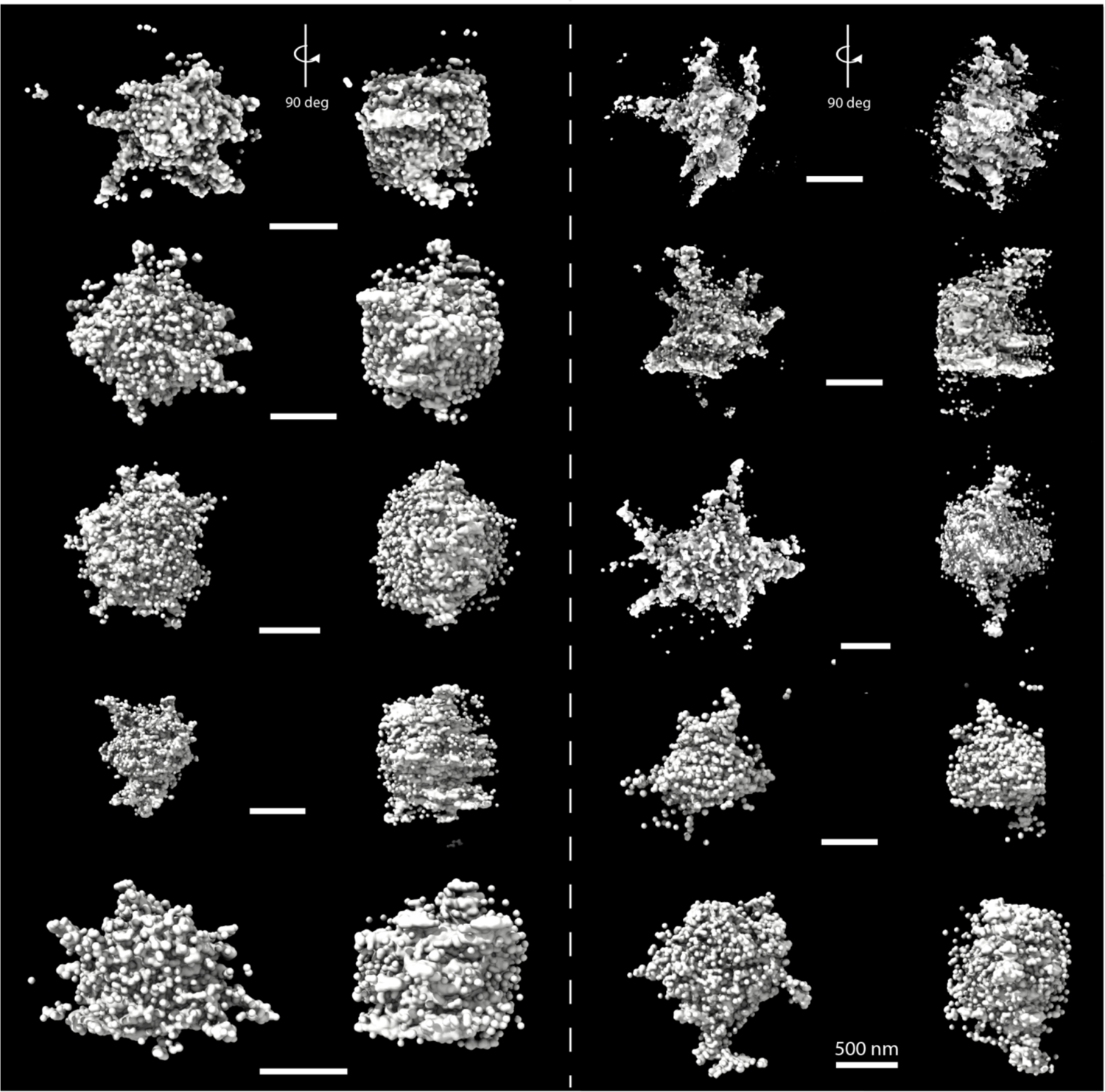
Three-dimensional super-resolution images of endogenous ASC specks. 3D renderings of ASC specks that were stained using an ASC antibody and a secondary Alexa Fluor 647-conjugated F(ab’)_2_ fragment and measured using dSTORM. Data was obtained on two independent cell preparations.

**Supplementary Figure S6:**
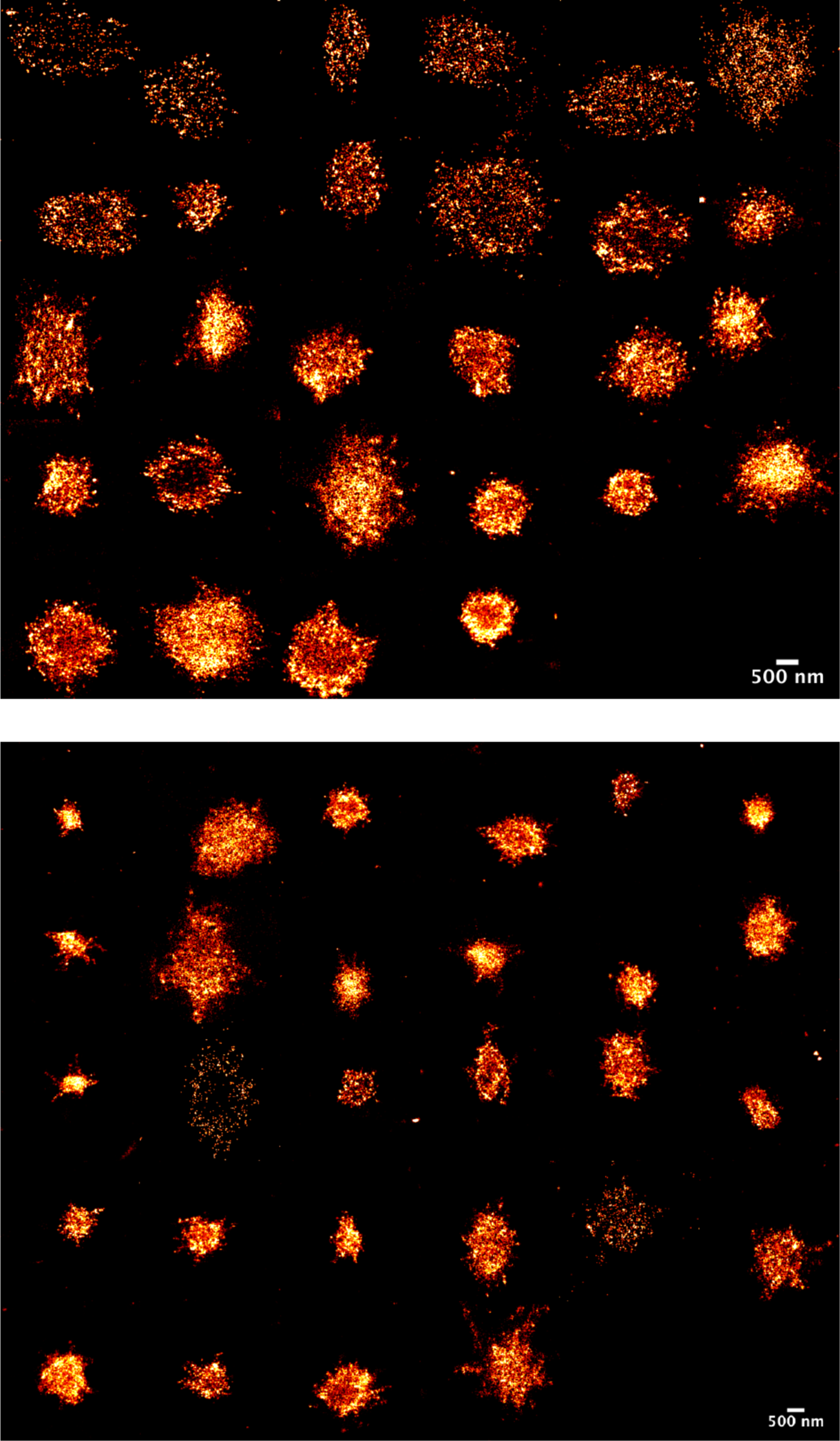
Two-dimensional renderings of 3D super-resolution images of endogenous ASC specks. The ASC specks were measured in 3D using dSTORM and stained with an ASC antibody and an Alexa Fluor 647-conjugated secondary F(ab’)_2_ fragment. Data was obtained on two independent cell preparations.

**Supplementary Figure S7:**
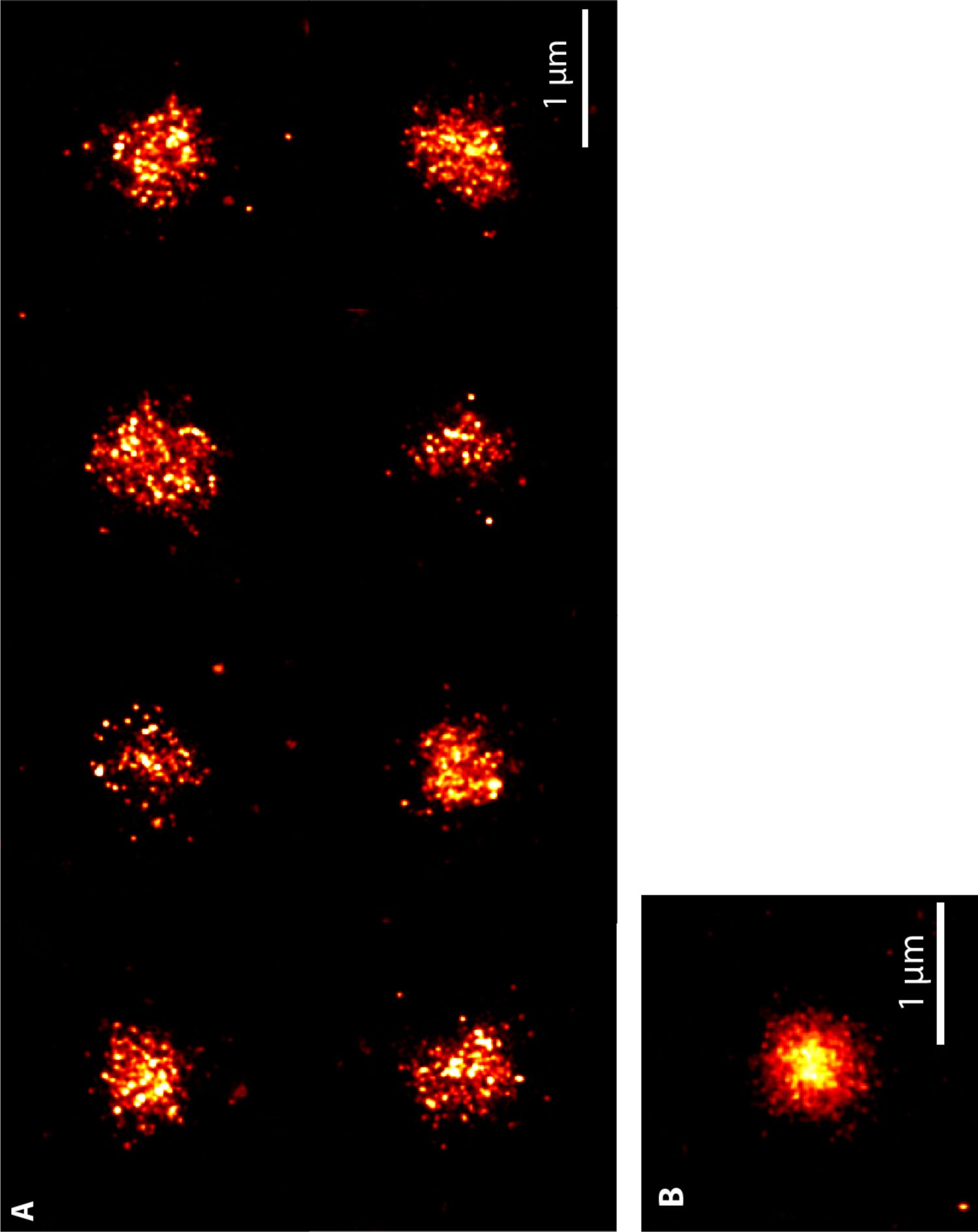
Nanobody labeling of endogenous ASC specks. A) ASC specks were stained with Alexa Fluor 647-conjugated nanobody and imaged using 2D STORM. Data was obtained on a single cell preparation. B) An ASC speck stained with a P3 DNA-PAINT docking strand-conjugated nanobody and imaged by 2D DNA-PAINT.

**Supplementary Figure S8:**
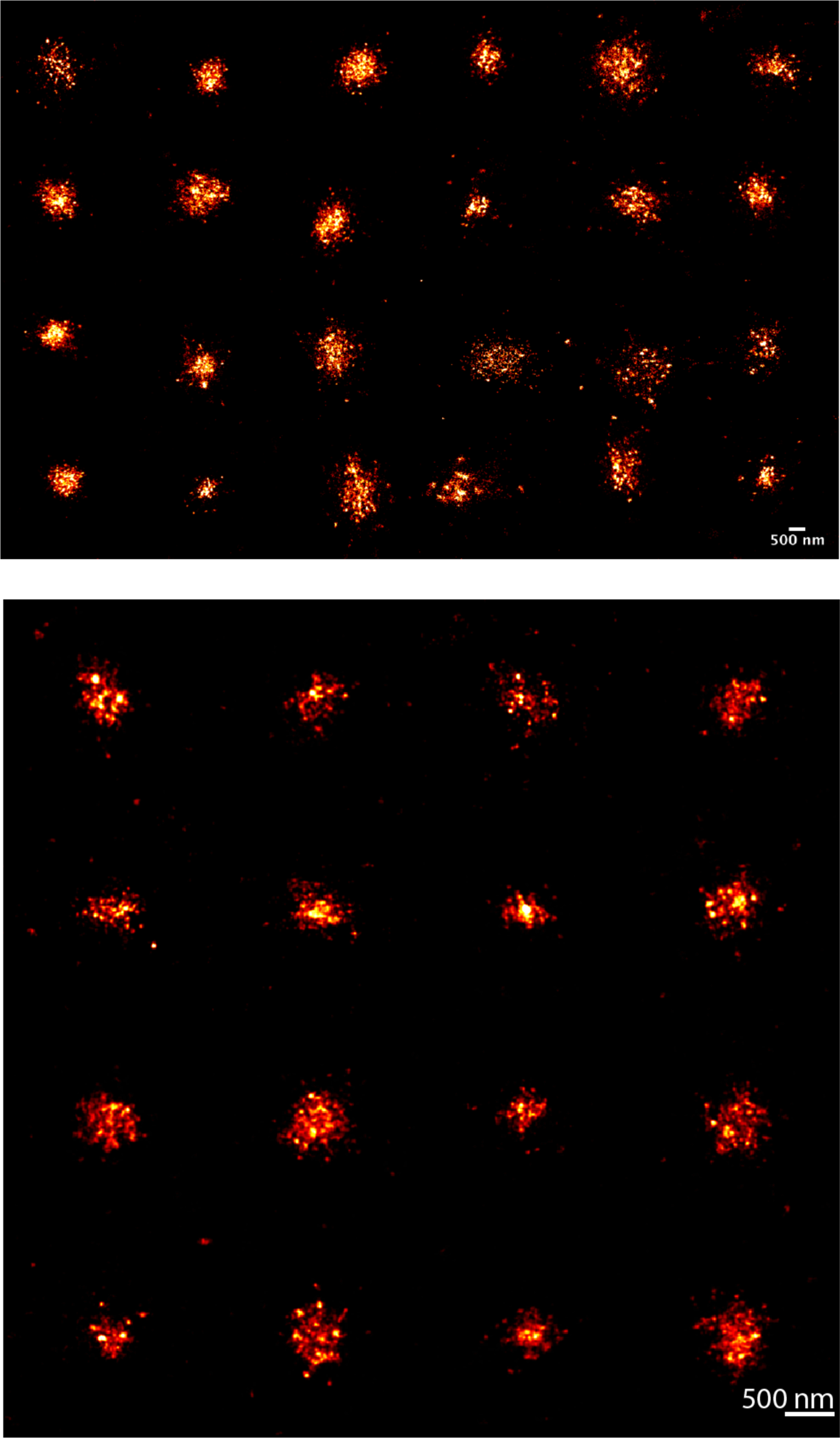
Two-dimensional renderings of nanobody-labeled endogenous ASC specks imaged in three-dimensions using dSTROM. The ASC nanobody was conjugated with Alexa Fluor 647.

**Supplementary Figure S9:**
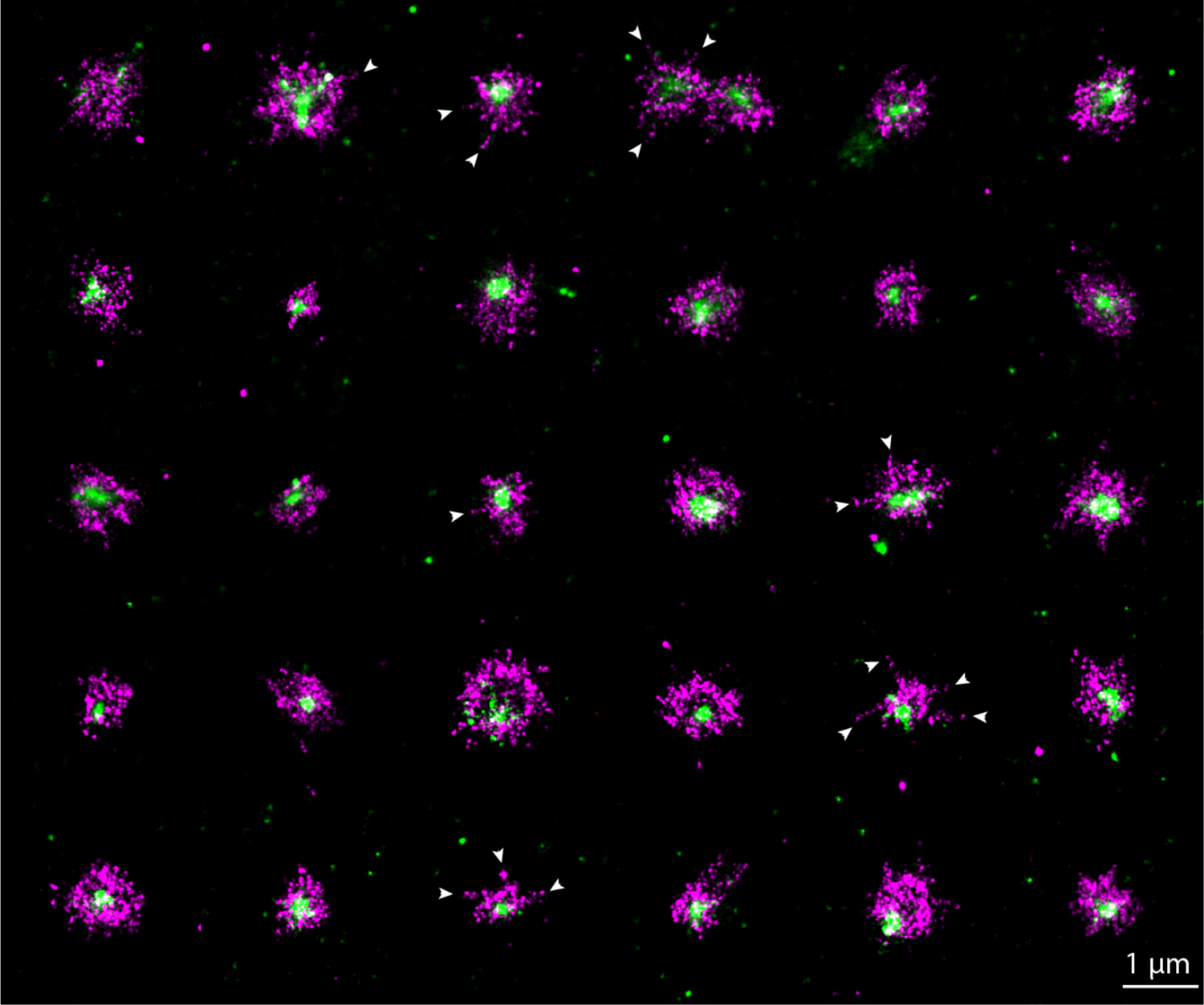
Two-color super-resolution images of endogenous ASC specks. 2D STORM images were collected of ASC specks labeled with primary ASC antibody and secondary Alexa Fluor 647-conjugated F(ab’)_2_ fragment (magenta), and with DyLight 755-conjugated ASC nanobody (green). Arrows indicate filamentous extensions reaching out from the speck center. Data was obtained on a single cell preparation.

**Supplementary Figure S10:**
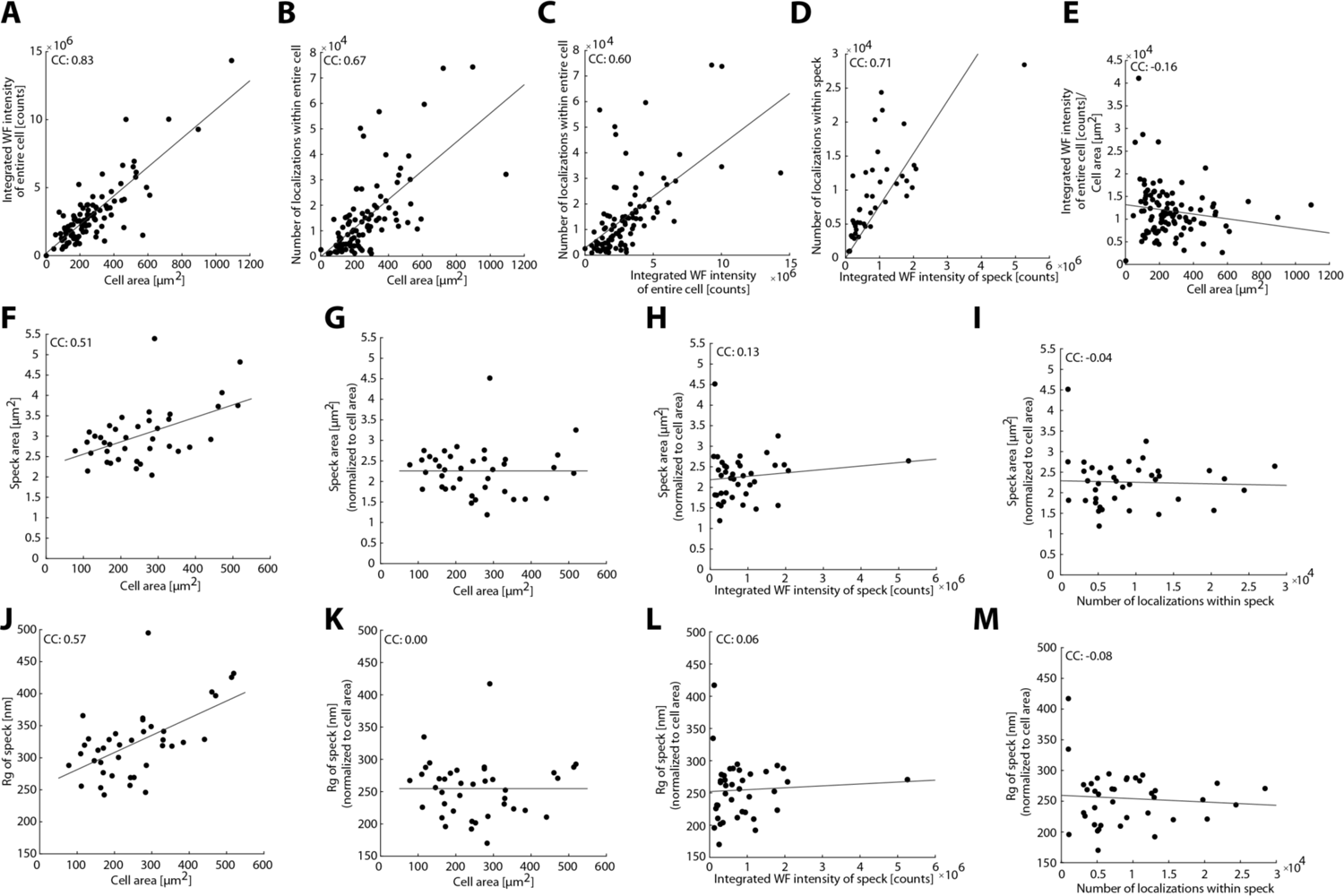
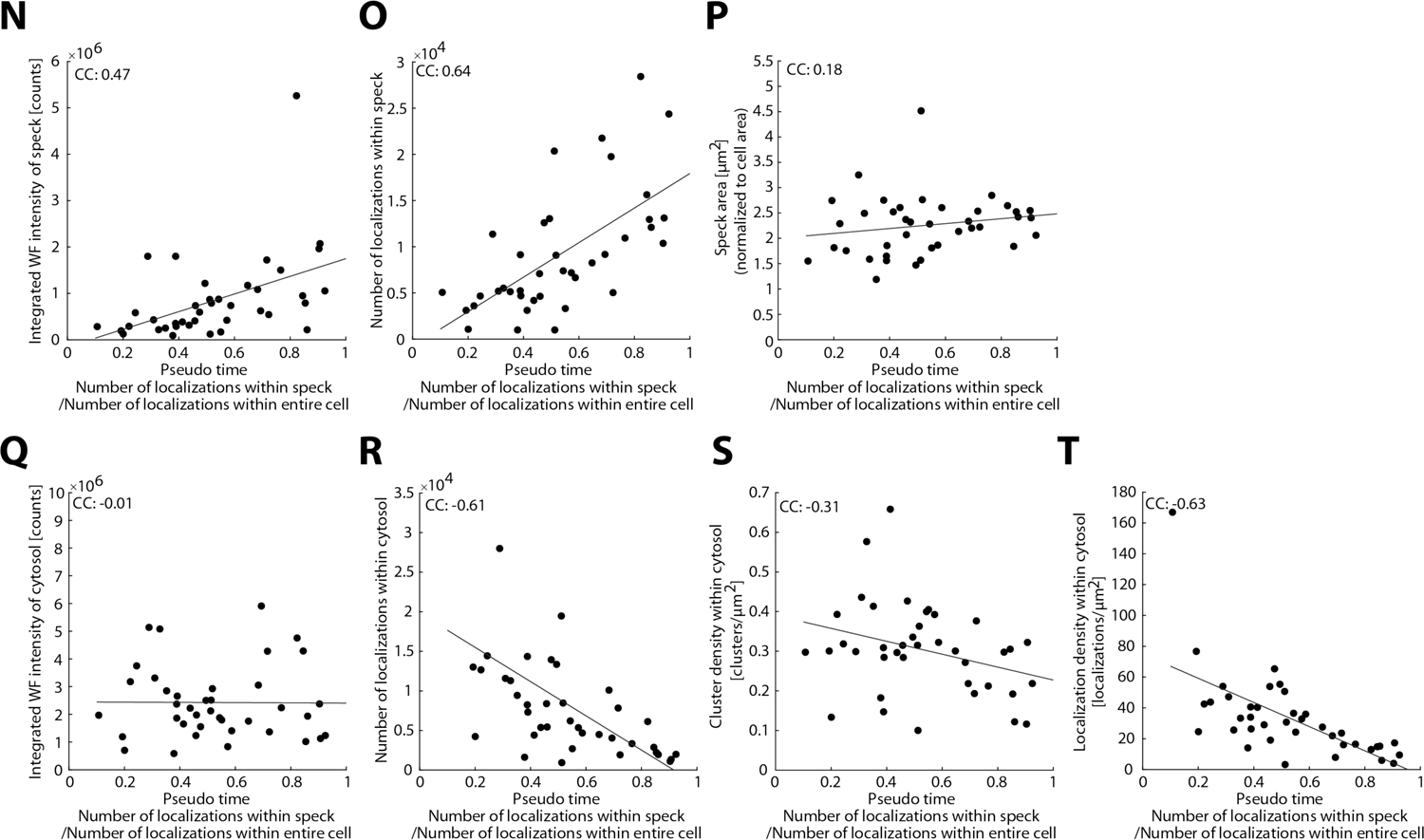
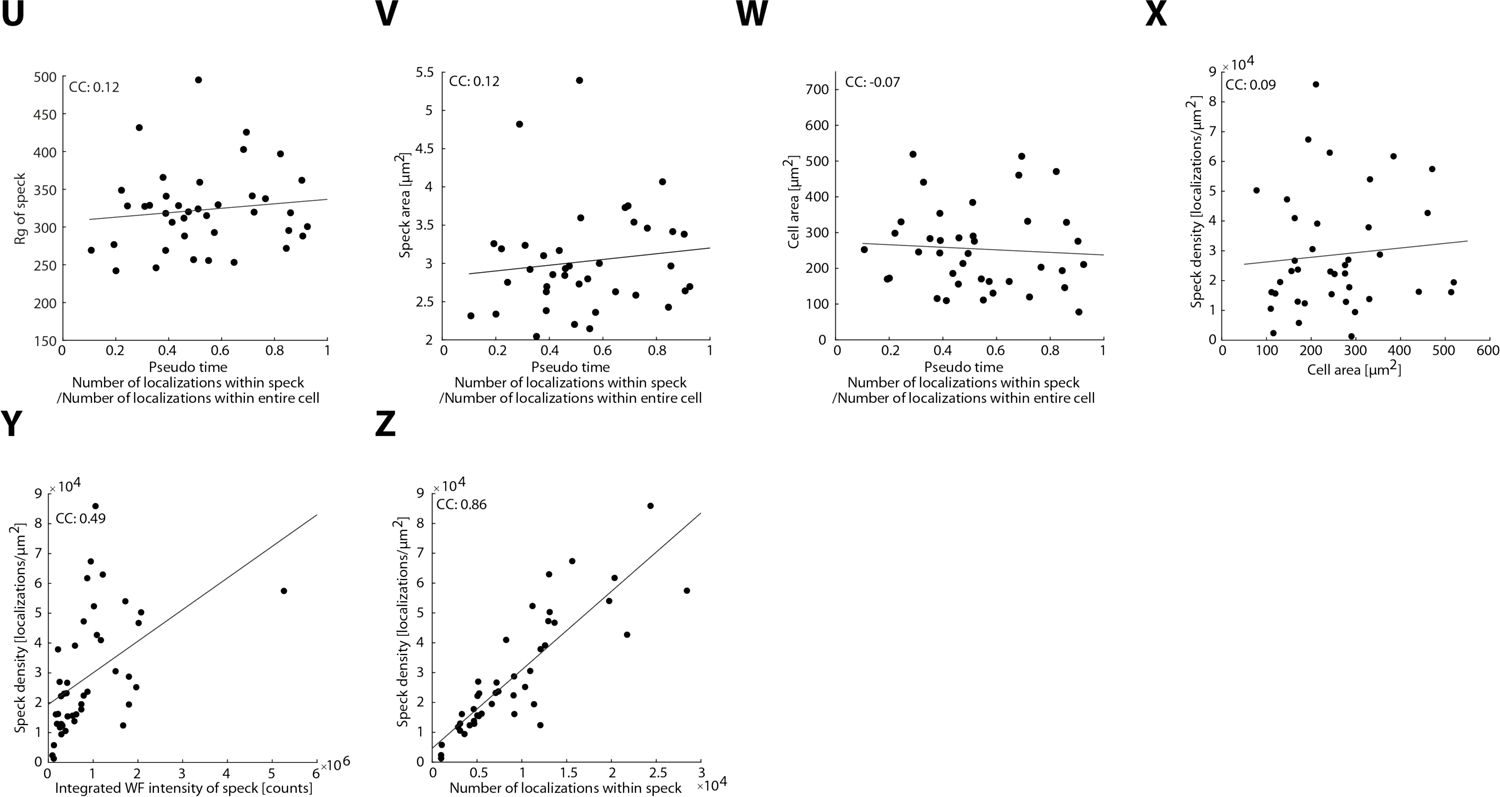
Correlation plots for various parameters obtained from the analysis of single cells. Panels A – E relate the number of localizations, the widefield intensity and the cell area with each other. Panels F – I depicts how the size of the speck depends on cell area and ASC concentration in the speck. Panels J - M depicts how the radius of gyration Rg, depends on cell area and ASC concentration in the speck. Panels N – X depict how various parameters depend on the pseudo time. Specifically, the amount of ASC in the speck (N, O), the speck area (P), the amount of ASC in the cytosol (Q, R), the cytosolic cluster density (S) and localization density (T), the radius of gyration, Rg (U), the speck area (V), the cell area (W) and the speck density (X). Panels Y – Z depict how the speck density depends on the amount of ASC in the speck. Data was obtained on three independent cell preparations. WF: widefield; CC: Pearson correlation coefficient

**Supplementary Figure S11:**
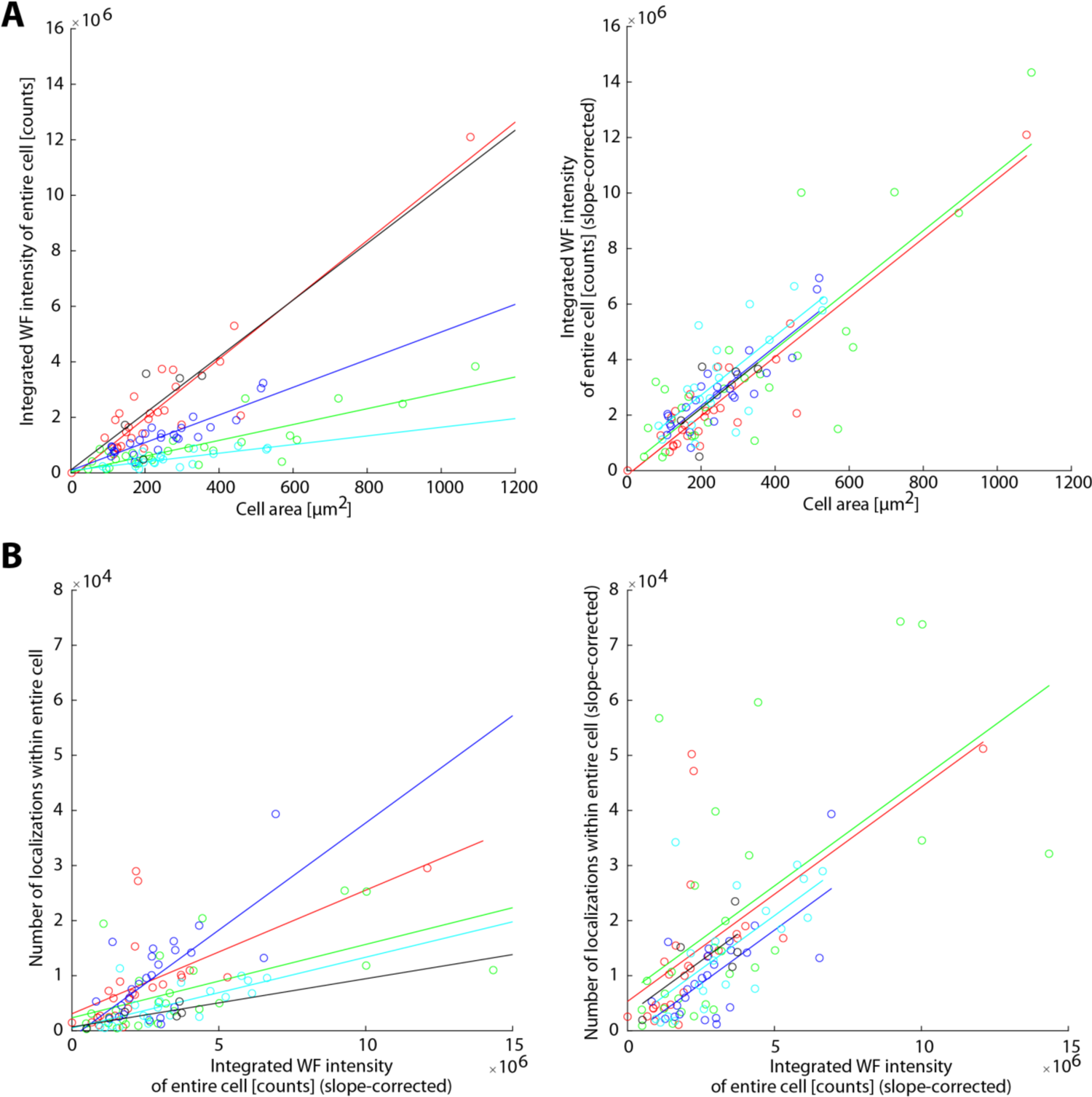
Scatter plots corrections. The uncorrected and corrected scatter plots are shown to illustrate the correction to the integrated widefield intensity and localization data for differences between measurement days. Different sample preparations and dSTORM buffer conditions lead to variations in the widefield intensity and number of localizations with respect to ASC content. Hence, we used the correlation between ASC content and cell area to normalize the widefield intensity for different measurement days. For the number of localizations, we used the linear correlation between intensity and number of detected localizations. A) Scatter plot of the integrated widefield intensity of the entire cell against the cell area including the linear fits of individual measurement days (left panel) and the normalized integrated widefield intensity by adjusting the individual slopes to that of the measurement day with the steepest slope against the cell area (right panel). B) Scatter plot of the number of localizations within the entire cell against the normalized integrated widefield intensity of the entire cell including the linear fits of individual measurement days (left panel) and number of localizations within the entire cell normalized with respect to the measurement day with the steepest slope plotted against the normalized integrated widefield intensity of the entire cell (right panel) The individual measurement days are color-coded.

**Supplementary Figure S12:**
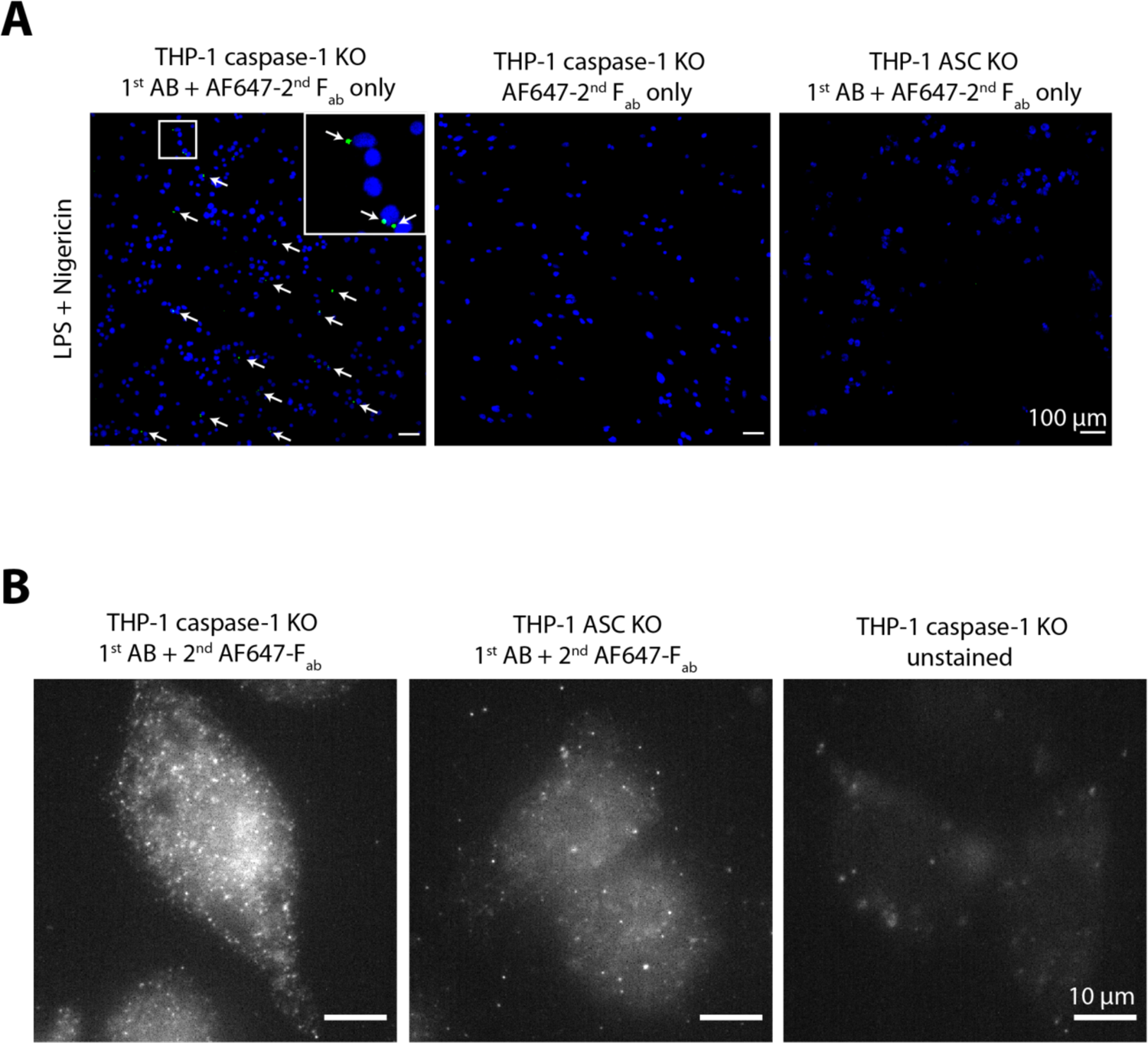
Controls for the applied antibody and F(ab’)*_2_* fragment staining in THP-1 knock-out (KO) cells. A) 10x overview images recorded on a confocal spinning disk microscope indicating the specificity of the applied ASC staining; green: Alexa Fluor 647 (summed projections of a recorded Z-stack are shown); blue: DNA-staining with DAPI. No ASC staining was observed in the absence of ASC or primary antibody. B) Higher magnification images (60x) verifying the specificity of ASC staining.

**Supplementary Figure S13:**
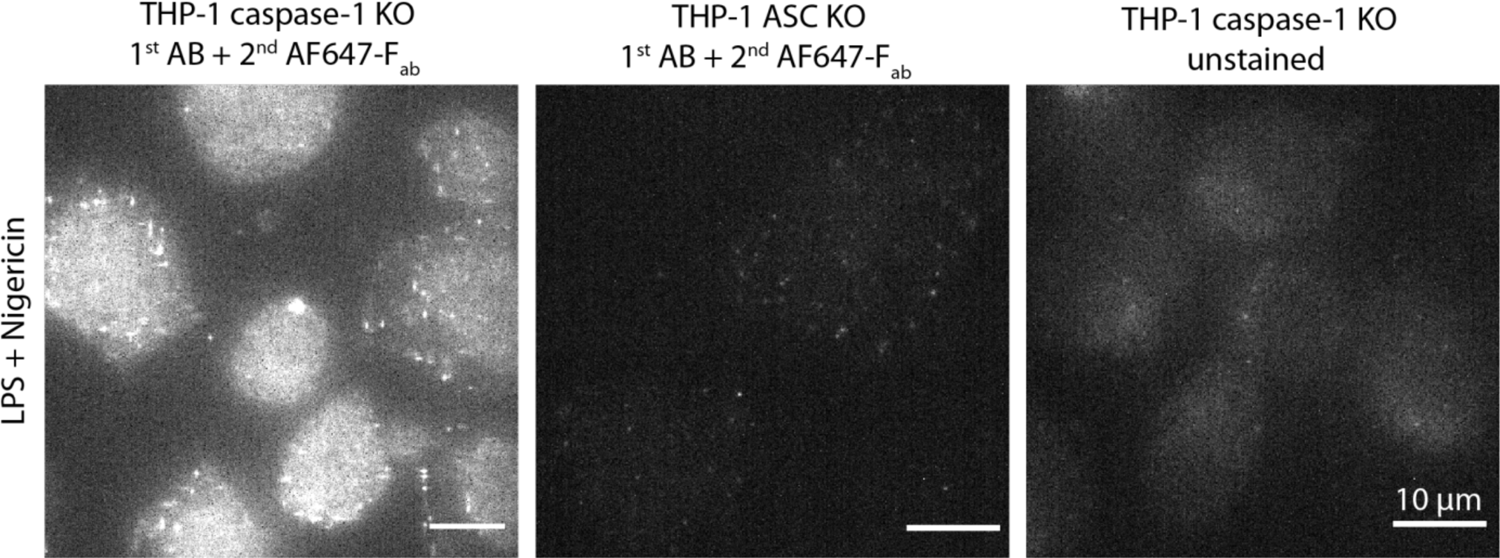
Controls for the applied nanobody (AF647-conjugated) staining in THP-1 knock-out (KO) cells. Widefield images recorded at 60x magnification are shown. Stimulated and labeled cells are shown in the left panel. No unspecific signal was observed in the absence of ASC (middle panel) or in unstained cells (right panel) verifying the specificity of ASC staining. All images are displayed using the same intensity scale.

## Captions Supplementary Movies

**Supplementary Movie S1: Three-dimensional super-resolution rendering of an endogenous ASC speck.** 3D rendering of an endogenous ASC speck stained with anti-ASC primary antibodies and secondary Alexa Fluor 647-conjugated F(ab’)_2_ fragments and measured using dSTORM. Note the overall spherical appearance characterized by a dense core and protruding filaments. The depicted complex exhibits a diameter of about 1 µm.

**Supplementary Movie S2: Three-dimensional super-resolution rendering of an endogenous ASC speck.** 3D rendering of an endogenous ASC speck stained with anti-ASC primary antibodies and secondary Alexa Fluor 647-conjugated F(ab’)_2_ fragments and measured using dSTORM. Note the overall spherical appearance characterized by a dense core and protruding filaments. The depicted complex exhibits a diameter of about 1 µm.

**Supplementary Movie S3: Three-dimensional super-resolution rendering of an endogenous ASC speck.** 3D rendering of an endogenous ASC speck stained with anti-ASC primary antibodies and secondary Alexa Fluor 647-conjugated F(ab’)_2_ fragments and measured using dSTORM. Note the overall ring-like appearance. The depicted complex exhibits a diameter of about 1 µm.

**Supplementary Movie S4: Three-dimensional super-resolution rendering of an endogenous ASC speck.** 3D rendering of an endogenous ASC speck stained with anti-ASC primary antibodies and secondary Alexa Fluor 647-conjugated F(ab’)_2_ fragments and measured using dSTORM. Note the overall ring-like appearance. The depicted complex exhibits a diameter of about 1 µm.

**Supplementary Movie S5: “Z-stack” super-resolution rendering of an endogenous ASC speck.** “Z-stack” 3D rendering of an endogenous ASC speck stained with anti-ASC primary antibodies and secondary Alexa Fluor 647-conjugated F(ab’)_2_ fragments and measured using dSTORM. Note the overall ring-like appearance.

**Supplementary Movie S6: “Z-stack” super-resolution rendering of an endogenous ASC speck.** “Z-stack” 3D rendering of an endogenous ASC speck stained with anti-ASC nanobodies conjugated to DNA-PAINT docking strands and measured using DNA-PAINT. Note the overall spherical appearance characterized by a dense core and protruding filaments.

